# α-ketoglutarate augments prolyl hydroxylase-2 mediated inactivation of phosphorylated-Akt to inhibit induced-thrombosis and inflammation

**DOI:** 10.1101/2021.06.11.448037

**Authors:** Nishith M Shrimali, Sakshi Agarwal, Simrandeep Kaur, Sulagna Bhattacharya, Sankar Bhattacharyya, Josef T Prchal, Prasenjit Guchhait

## Abstract

Phosphorylation of Akt (pAkt) regulates multiple physiological and pathological processes including thrombosis and inflammation. In an approach to inhibit the pathological signalling of pAkt by prolyl-hydroxylase-2 (PHD2) we employed α-ketoglutarate (αKG), a cofactor of PHD2. Octyl-αKG supplementation to platelets promoted PHD2 activity through elevated intracellular αKG:succinate ratio and reduced aggregation *in vitro* by suppressing pAkt1(Thr308). Augmented PHD2 activity was confirmed by increased hydroxylated-proline alongside enhanced binding of PHD2 to pAkt in αKG-treated platelets. Contrastingly, inhibitors of PHD2 significantly increased pAkt1 in platelets. Octyl-αKG followed similar mechanism in monocytes to inhibit cytokine secretion *in vitro*. Our data also describe a suppressed pAkt1 and reduced activation of platelet and leukocyte obtained from mice supplemented with dietary-αKG, unaccompanied by alteration in their counts. Dietary-αKG significantly reduced clot formation and leukocyte accumulation in various organs including lung of mice treated with thrombosis-inducing agent carrageenan. Importantly, we observed a significant rescue effect of dietary-αKG on inflamed lung of SARS-CoV-2 infected hamsters. αKG significantly reduced leukocyte accumulation, clot formation and viral load alongside downmodulation of pAkt in lung of the infected animals. Therefore, our study suggests a safe implementation of dietary-αKG in prevention of Akt-driven anomalies including thrombosis and inflammation, highlighting a better pulmonary management in COVID-19.

## Introduction

The serine-threonine kinase Akt, also known as protein kinase B (PKB), contributes to a broad range of cellular functions including cell survival, proliferation, gene expression and migration of cells of most lineages. Akt plays a central role in both physiological and pathological signalling mechanisms. Upon exposure to stimuli, Akt is recruited to the cell membrane by phosphoinositide 3-kinase (PI3K), where it is phosphorylated by membrane associated 3-phosphoinositide-dependent kinase-1 (PDK1) and therefore activated. Among the three isoforms, Akt1 is widely expressed in most of the cell types in both human and mice (1-5). The poignant role of PI3K-Akt signalling is well investigated in platelet activation and functions including aggregation, adhesion and thrombus formation (1-7). Platelets from Akt1-/- mice displayed an increased bleeding time and in *ex vivo* their platelets minimally responded to agonists (3). Studies using inhibitors to Akt including SH-6, triciribine and Akti-X describe the important role of this signaling adaptor molecule in platelets functions including aggregation, clot formation and granule secretion *in vitro* and *in vivo* (4, 8-9).

Extensive studies have reported the crucial involvement of the PI3K-Akt pathway in the regulation of immune cell function in a broad range of inflammatory diseases such as rheumatoid arthritis, multiple sclerosis, asthma, chronic obstructive pulmonary disease, psoriasis, and atherosclerosis (10-15). Besides, the pathological signalling of Akt is well reported in the progression cancer or tumor cells (16). Activation of PI3K-Akt pathway is highly relevant in infections like SARS-CoV-2 (17-18), SARS-CoV (19), Dengue and Japanese Encephalitis (20) viruses. Akt signalling has been found to be pivotal for the virus entry and replication in host cells. Therefore, extensive reports suggest Akt as a potential therapeutic target in various disease conditions (21). The PI3K-Akt inhibitor wortmannin has been used to alleviate the severity of inflammation and improve the survival rate in rats with induced severe acute pancreatitis. Akt1-/- mice showed a markedly reduced carrageenan-induced paw edema and related inflammation alongside a significant decrease in neutrophil and monocyte infiltration (22). Studies using inhibitors such as Akti-8 and Akt-siRNA describe the crucial regulatory role of Akt in inflammatory response of monocytes and macrophages *in vitro* and *in vivo* (23-24).

In the context of regulation of the Akt pathway, a study has reported that phosphorylated Akt1 (pAkt1) is hydroxylated by an oxygen-dependent enzyme, prolyl hydroxylase 2 (PHD2). The pAkt1 undergoes prolyl hydroxylation at Pro125 and Pro313 by PHD2 in a reaction decarboxylating α-ketoglutarate (αKG). This promotes von Hippel–Lindau protein (pVHL) binding to the hydroxylated site. pVHL then interacts with protein phosphatase 2A (PP2A), which dephosphorylates the Thr308 site, resulting in Akt1 inactivation (25).

In this study, we describe a heretofore unreported role of PHD2 in the regulation of platelet and monocyte functions by inactivating pAkt1. Supplementation with dietary αKG, a metabolite of TCA cycle and a cofactor of PHD2, appears to be a potent suppressor of pAkt, significantly reducing clot formation and leukocyte accumulation and related thrombotic and inflammatory events in mice treated with thrombosis-inducing agent like carrageenan (26). Further, we show a rescue effect of αKG on lung inflammation in SARS-CoV-2 infected hamsters. SARS-CoV-2 infection to golden hamsters induces significant inflammation of the bronchial epithelial cells and lung, in turn increases the disease severity between days 3-5 (27). SARS-CoV-2 significantly increases phosphorylation of Akt1(Thr308) (17), a known target of PHD2, in infected cells. We describe that dietary αKG significantly reduced clot formation, inflammation and viral load in conjunction with downmodulation of pAkt in the lungs of hamsters with SARS-CoV-2 infection.

## Results

### Agonist mediated platelet activation is directly related to phosphorylation of Akt1 but inversely with prolyl-hydroxylase activity of PHD2

We checked the presence of all 3 isoforms PHD1, PHD2 and PHD3 in platelets (Figure 1A). A recent report has shown that PHD2 hydroxylates the proline residues of phosphorylated Akt1(Thr308) [pAkt1(Thr308)] eventually leading to inactivation (25). Therefore, we measured phosphorylation status of Akt1 in agonist-activated platelets and check if the PHD2 activity is functionally relevant. As expected, the phosphorylation of Pan-Akt or Akt1(Thr308) was increased with higher concentrations of agonists like collagen (Figure 1B-D) and ADP (Supplemental Figure 1A), indicating an insufficient activity of PHD2 enabling the hydroxylation of elevated level of pAkt after agonist stimulation. To confirm the prolyl-hydroxylation of elevated level of pAkt after agonist stimulation. To confirm prolyl-hydroxylation activity of PHD2, we measured expression of other known substrates of the enzyme, such as HIF1α and HIF2α in agonist-activated platelets. We observed stabilization of both HIF1α and HIF2α in activated platelets under normal oxygen condition or normoxia (Figure 1B). It pertinently raised the speculation that the enzymatic activity of PHD2 remains below the threshold level to inactivate pAkt, which heightens in platelets after agonist stimulation. We then tested whether agonist or other chemical induced alteration in prolyl-hydroxylase activity of PHD2 can in turn alter the activation status of pAkt in platelet?

**Figure 1.**
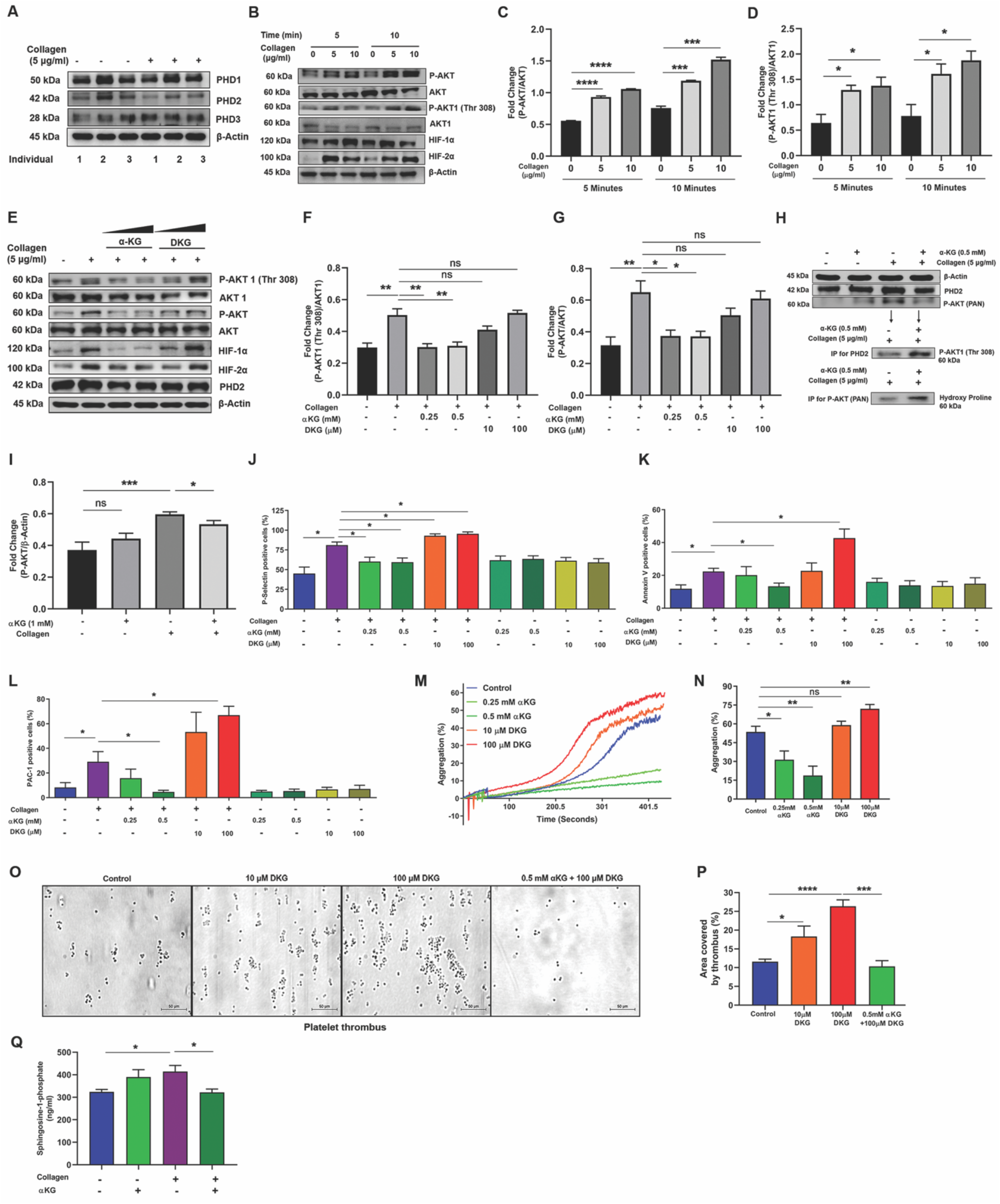
*α*KG mediated augmentation of prolyl-hydroxylase activity of PHD2 inactivate pAkt in platelets. (**A**) All 3 isoforms of PHD exist in human platelet. Platelet-rich plasma (PRP) isolated from 3 different healthy individuals, after adjusting to equal number of platelets, was incubated with or without collagen (5 µg/ml) for 5 min and processed for western blotting (WB) of PHD1, PHD2 and PHD3. Densitometry data are mentioned in Supplemental Figure 14A. (**B-G**) PRP was incubated with collagen in a time- and concentration-dependent manner, and (B) expression of phosphorylated (p)-Akt, Akt, pAkt1(Thr 308), Akt1, HIF-1α and HIF-2α was measured in platelet pellet using WB. (C-D) Densitometry analysis shows elevated expression of pAkt and pAkt1 after collagen stimulation, other densitometry data are mentioned in Supplemental Figure 14B-C. (E) Platelets were incubated with collagen in presence of αKG (0.25 and 0.5 mM) or DKG (10 and 100 µM) and the expression of above signaling molecules and PHD2 was measured using WB. (F-G) Densitometry data show suppression of collagen-induced elevation of pAkt and pAkt1 by αKG, but an elevation of these molecules in presence of DKG. Other densitometry data are mentioned in Supplemental Figure 14D-E. (**H-I**) (H) Immunoprecipitation (IP) of PHD2 from lysate of washed-platelets from above experiment was performed and processed for WB of pAkt; further, IP of pAkt from same lysate and WB for hydroxy proline shows the interaction between the molecules. (I) Densitometry data of pAkt. Other densitometry data are mentioned in Supplemental Figure 14F-G. (**J-L**) PRP from above experiment of Figure 1C was processed for measuring surface (J) P-Selectin, (K) PS (Annexin-V binding) and (L) GPIIbIIIa activation (PAC-1 binding) using flow cytometry. (**M-N**) (M) Platelet aggregation was performed using PRP pre-treated with αKG or DKG in response to collagen. (N) Percentage platelet aggregation was measured. Data from similar experiment in response to agonist ADP are mentioned in Supplemental Figure 4. (**O-P**) (O) PRP from healthy individuals was incubated with αKG or DKG and perfused on immobilized collagen surface under arterial flow share condition 25 dyne/cm^2^ and platelet thrombus formation was measured. Scale bar 50 µm. (P) Thrombus area was measured. (**Q**) Secretion of Sphingosine-1-phosphate (S1P) was quantified from supernatant of αKG- and collagen-treated washed platelets using ELISA. Data in above figure are Mean ± SEM from 3 independent experiments. Unpaired t-test was used to compare between the groups, \**P*<0.05. \*\**P*<0.01, \*\*\**P*<0.001, \*\*\*\**P*<0.0001 and ns=non-significant.

### Supplementation of octyl αKG inhibits Akt1 phosphorylation by augmenting PHD2 activity

To alter the enzymatic activity of PHD2, we used α-ketoglutarate (αKG, a cofactor of PHD2) and dimethyl ketoglutarate (DKG, an inhibitor to PHD2). Octyl αKG (a membrane-permeating form) supplementation significantly decreased the collagen-induced phosphorylation of Akt1(Thr308) or pan-Akt, and also destabilized HIF1α and HIF2α in platelets under normoxia (Figure 1E-G). Interaction of pAkt1 with PHD2 was confirmed by immunoprecipitation of PHD2 followed by immunoblotting for pAkt1(Thr308). A significantly increased binding of PHD2 to pAkt1(Thr308) in presence of αKG, suggests a pre-existing condition of elevated prolyl-hydroxylation and inactivation of pAkt1 in collagen-activated platelets (Figure 1H). In order to check the enzymatic activity of PHD2 on pAkt, immunoprecipitation was performed using pan-pAkt antibody and immunoblotted for hydroxy proline. An increased level of pAkt-bound hydroxy proline (Figure 1H), suggests an elevated hydroxylation of proline on pAkt in platelets after αKG supplementation. On the other hand, the PHD2 inhibitor DKG promoted the Akt phosphorylation (Figure 1E-G). Other known inhibitor of PHD2, such as ethyl-3-4-dihydroxybenzoic acid (DHB) also exhibited an elevated Akt phosphorylation (Supplemental Figure 1B). Although, αKG and DKG altered the enzymatic function of the PHD2 to regulate pAkt, neither of the treatments altered the expression of PHD2 in platelets significantly (Figure 1E). We examined the effect of αKG on PI3K, activator of Akt. Our data show no significant effect of αKG on the expression of phosphorylated PI3K(p55) (Supplemental Figure 1C), thus, confirming its specific target on pAkt.

### Octyl αKG supplementation suppresses agonist-induced activation, aggregation and thrombus formation of platelet while PHD2 inhibitors enhance it

Octyl αKG supplementation suppressed the expression of cell surface activation markers such as P-selectin, phosphatidylserine (PS) and PAC-1 binding to GPIIbIIIa integrin on collagen-activated platelets (Figure 1J-L) and microparticle release from activated platelets (Supplemental Figure 3) in a concentration-dependent manner *in vitro*. In contrast, DKG enhanced the above parameters (Figure 1J-L). Similarly, αKG suppressed platelet aggregation induced by collagen (Figure 1M-N) or ADP (Supplemental Figure 4A-B) in a concentration-dependent manner, but DKG enhanced it. Another PHD2 inhibitor DHB also enhanced collagen-induced platelet aggregation (Supplemental Figure 4C-D). Further, our data show that platelet thrombus formation was increased in a dose-dependent manner when whole blood was treated with DKG and perfused under flow shear condition on immobilized collagen surface. αKG significantly suppressed DKG-induced thrombus formation (Figure 1O-P). Collagen-activated platelets secreted large amount of sphingosine-1-phosphate (S1P), known stimulator of monocytes, which too was reduced by αKG supplementation (Figure 1Q).

### Octyl αKG supplementation significantly suppresses monocyte functions by augmenting PHD2 activity

We then investigated the PHD2-mediated inhibition of pAkt1(Thr308) in monocytes, isolated from healthy individuals and activated with either S1P or LPS after pre-treatment with octyl αKG *in vitro*. Our data show that the αKG supplementation decreased pAkt1(Thr308) and increased degradation of HIF2α (Figure 2A-C). Simultaneous suppression in secretion of inflammatory cytokines including, IL1β, IL6, TNFα and IL10 was observed (Figure 2D-G). Similar outcomes were observed in monocytes exposed to LPS after pre-treatment of αKG (Supplemental Figure 5). We then confirmed that the above mechanism of αKG-induced suppression of monocyte activation is mediated primarily by pAkt, independent of HIFα. In HIF1α-depleted U937 monocytic cells (detailed protocol of shRNA-mediated depletion is mentioned in Supplemental Figure 6), αKG supplementation significantly reduced cytokine secretion (Figure 2H-I) alongside down-modulated pAkt1(Thr308) (Figure 2J-K).

**Figure 2.**
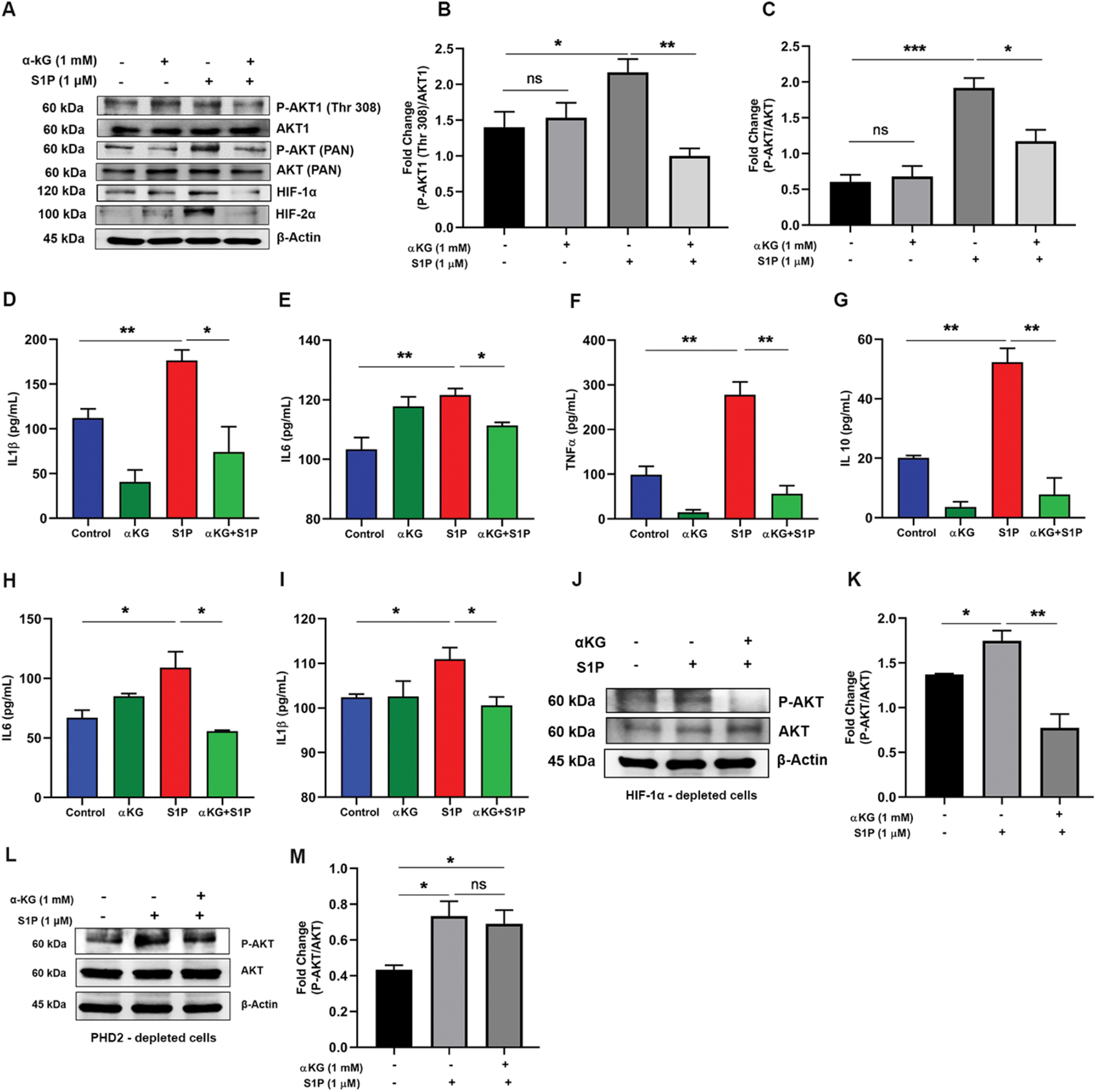
*α*KG mediated elevation in prolyl-hydroxylase activity of PHD2 inhibits Akt phosphorylation in monocytes. (**A-C**) Monocytes isolated from healthy individuals were activated with S1P with/without pre-treatment with αKG. (A) The expression of pAkt, Akt, pAkt1(Thr 308), Akt1, HIF-1α and HIF2α was measured using WB. (B-C) Densitometry data show suppression of S1P-induced elevation of pAkt and pAkt1 by αKG. Other densitometry data are mentioned in Supplemental Figure 14I-J. Data from similar experiment using LPS as an agonist to monocytes, are mentioned in Supplemental Figure 5. (**D-G**) The cytokines such as (D) IL1β, (E) IL6, (F) TNFα and (G) IL10, secreted by monocytes from above experiment, were measured using flow cytometry based CBA array. (**H-K**) U937 monocytic cell line was depleted for *HIFA* (HIF1α) using shRNA (detailed protocol is mentioned in Supplemental Figure 6) and treated with αKG before exposing to S1P treatment. (H) IL6 and (I) IL1β were measured from supernatant. (J-K) The cells from above experiment were used for measuring expression of pAkt and Akt using WB. (**L-M**) PHD2-depleted U937 monocytic cells were treated with αKG before exposing to S1P treatment and used for measuring expression of pAkt and Akt using WB. Data in above figure are Mean ± SEM from 3 independent experiments. Unpaired t-test was used to compare between the groups, \**P*<0.05, \*\**P*<0.01 and ns= non-significant.

### αKG-mediated suppression of pAkt in monocyte is mediated by PHD2

To confirm that αKG-mediated suppression of pAkt in monocyte is mediated by PHD2, we performed the above experiment in PHD2-depleted U937 monocytic cells. Our data show that S1P-induced elevation of pAkt1 was not significantly suppressed by octyl αKG in PHD2-depleted cells (Figure 2L-M), highlighting the role of PHD2 in the inactivation of pAkt1.

### Elevated intracellular ratio of αKG to succinate correlates with augmented activity of PHD2 in platelet and monocyte after octyl αKG supplementation

To ascertain the mechanism of augmentation of PHD2 activity we measured an elevated level of intracellular αKG in collagen-activated platelets after octyl αKG supplementation, although the level was unaltered in collagen-activated platelets compared to resting platelets *in vitro* (Figure 3A). Since, succinate, a product of αKG-dependent dioxygenase reaction in TCA cycle, inhibits PHD2 function; we measured its intracellular level and found no significant change in platelets after collagen activation as well as post αKG supplementation (Figure 3B). However, the intracellular αKG to succinate ratio was elevated in collagen-activated platelets after αKG supplementation, which might have played a role in augmentation of PHD2 activity (Figure 3C) as suggested by others (28) as well as our recent work (29). The intracellular level of other metabolites such as fumarate and pyruvate were found unaltered (Supplemental Figure 7A-B). We observed elevated level of lactate in the supernatant of activated platelets, which was reduced by αKG supplementation (Figure 3D).

**Figure 3.**
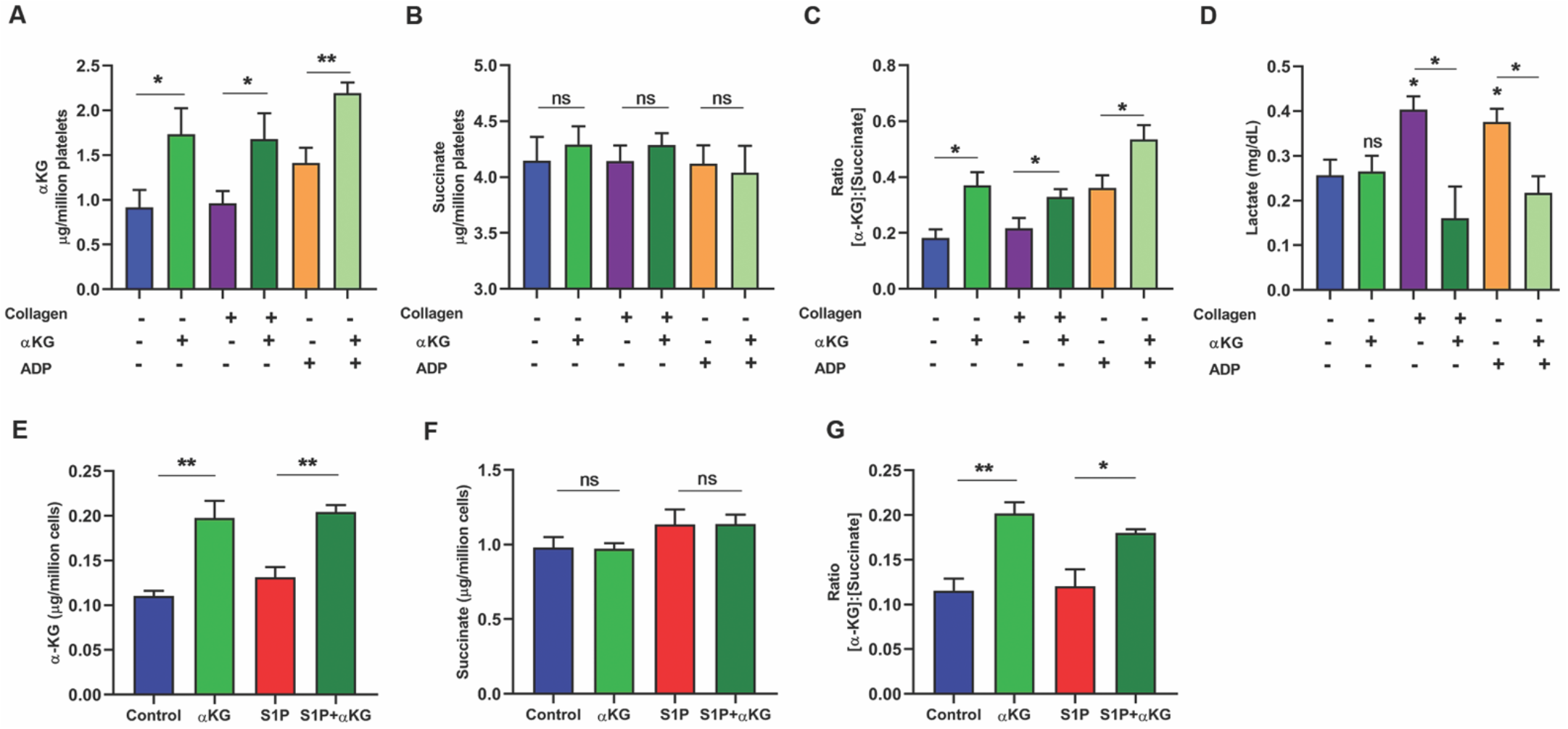
Elevation in intracellular ratio of *α*KG to succinate promotes PHD2 activity in both platelet and monocyte. (**A-D**) PRP isolated from healthy individuals was pre-treated with αKG and activated in presence of collagen/ADP. Platelet pellet was washed and lysed, and used for measuring intracellular (A) αKG and (B) succinate using colorimetry based assay. (C) Represents intracellular ratio of αKG to succinate. (D) Release of Lactate from supernatant of platelets from above experiment was quantified. Data of Pyruvate and Fumarate from platelet lysate is mentioned in Supplemental Figure 7. (**E-F**) Similarly, intracellular (E) αKG and (F) succinate were measured from cell lysate of primary monocytes pretreated with αKG and stimulated with S1P as mentioned in Figure 2A. (G) Represents ratio of αKG to succinate in monocytes. Data in above figure are Mean ± SEM from 3 independent experiments in duplicates. Unpaired t-test was used to compare between the groups, \**P*<0.05 and \*\**P*<0.01, ns=non-significant.

We observed a similar elevation of intracellular αKG to succinate ratio in S1P-stimulated monocytes after octyl αKG supplementation, although the ratio was unaltered in SIP-activated monocytes compared to untreated monocytes (Figure 3 E-G), which might have played a role in augmentation of PHD2 activity.

### Supplementation of dietary αKG significantly inhibits platelet aggregation in mice

We investigated whether the supplementation of dietary αKG inhibits platelet aggregation in mice, and observed that 1% αKG via drinking water for 24 and 48 hrs (experimental detailed is mentioned in Figure 4A) significantly inhibited platelet aggregation *ex vivo* in response to agonists such as collagen (Figure 4B-C) and ADP (Supplemental Figure 9C). The above αKG supplementation did not alter the counts of platelets and WBCs in mice (Supplemental Figure 8A-C), suggesting a safe implementation of the metabolite. In a recent work, we have mentioned the safe rescue effect of 1% dietary αKG in mice exposed to hypoxia treatment (29).

**Figure 4.**
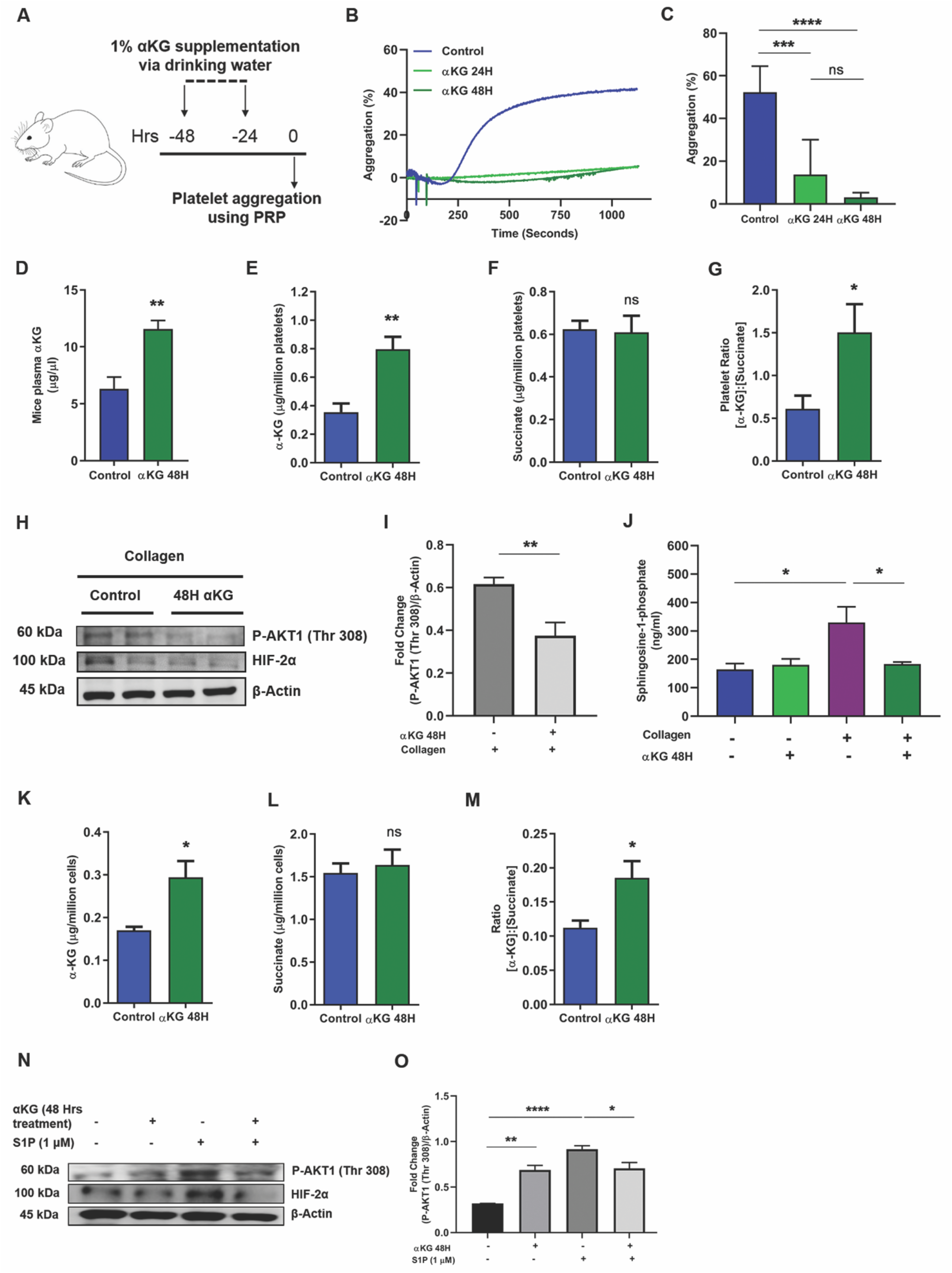
Dietary *α*KG supplementation to healthy wild type mice reduces platelet and monocyte activation. (**A**) Schematic showing experimental protocol. BALB/c mice were supplemented with dietary αKG for 24 and 48 hrs, and whole blood was isolated to separate PRP for assessing following parameters. (**B-C**) (B) Percentage platelet aggregation in presence of collagen or ADP (ADP data in Supplemental Figure 9). (C) Percentage platelet aggregation from 8 mice in each group. (**D-G**) (D) Plasma concentration of αKG was quantified from above mice, control vs. 48 hrs αKG groups (n=8). The intracellular (E) αKG and (F) succinate and their (G) fold change ratio in platelets of mice pre-treated with/without αKG for 48 hours (n=8). (**H-I**) (H) Expression of pAkt1(Thr 308), HIF-2α was measured in platelets from mice of above groups using WB. Representative image shows blot from 2 mice. (I) Densitometry data show suppression of collagen-induced elevation of pAkt1(Thr 308) in αKG treated mice (n=8). Other densitometry data in Supplemental Figure 14K. (**J**) S1P secretion was measure from supernatant of collagen-activated platelets of above mice groups (n=8). (**K-M**) Similarly, the intracellular (K) αKG and 1.succinate and their (M) fold change ratio in PBMCs was detected from above mice (n=8). (**N-O**) (N) Expression of pAkt1(Thr 308), HIF-2α was measured in PBMCs from above mice, after treating the cells with S1P *ex vivo*. Representative image shows blot from a mouse. (O) Densitometry data show suppression of S1P-induced elevation of pAkt in αKG treated mice (n=8). Other densitometry data in Supplemental Figure 14L. Data in above figure are Mean ± SEM from 3 independent experiments. Unpaired t-test was used to compare between the groups, \**P*<0.05, \*\**P*<0.01, \*\*\**P*<0.001, **** *P*<0.0001 and ns=non-significant.

### Elevated intracellular ratio of αKG to succinate relates augmented activity of PHD2 in platelet and monocyte of mice with dietary αKG supplementation

We have investigated the enzymatic activity of PHD2 in mice from above experiment. Our data show that the dietary αKG supplementation elevated αKG level in plasma (Figure 4D) and also in platelets (Figure 4E). An increased intracellular ratio of αKG to succinate in platelets (Figure 4F-G) might have augmented PHD2 activity. In fact, the decreased expression of pAkt1 and HIF2α in platelets of αKG-supplemented mice, confirmed the augmented-activity of PHD2 (Figure 4H-I). The platelets from αKG-supplemented mice showed a reduced secretion of inflammatory mediator S1P (Figure 4J). Similarly, PBMCs collected from αKG-supplemented mice displayed elevated level of intracellular αKG as well as αKG to succinate ratio (Figure 4K-M) along with downmodulation of pAkt and HIF2α (Figure 4N-O). We then tested the effect of dietary αKG supplementation in animal models of induced thrombosis and inflammation.

### Dietary αKG significantly inhibits carrageenan-induced thrombosis and inflammation in mice

Dietary supplementation of αKG (starting at 24 or 48 hrs before carrageenan treatment) significantly reduced the tail thrombosis (Figure 5B-C) and clot formation in lungs and liver (Figure 5D-G, Supplemental Figure 10A-B), as well as, accumulation of leukocytes in lungs (Figure 5H-I, Supplemental Figure 11) of mice exposed to carrageenan intraperitoneal treatment for 48 hrs. αKG supplementation also suppressed the levels of inflammatory cytokines such as IL1ß, IL6 and TNFα in plasma of the carrageenan treated mice (Figure 5J-M). Besides, αKG-supplemented mice when treated with carrageenan locally at abdomen and for a short period (3-6 hrs), displayed less accumulation of leukocytes including monocytes and neutrophils, and leukocyte-platelet aggregates, as well as, decreased pro-inflammatory agents such as myeloperoxidase (MPO) in the peritoneum (Figure 5N-T; Supplemental Figure 12B-C).

**Figure 5.**
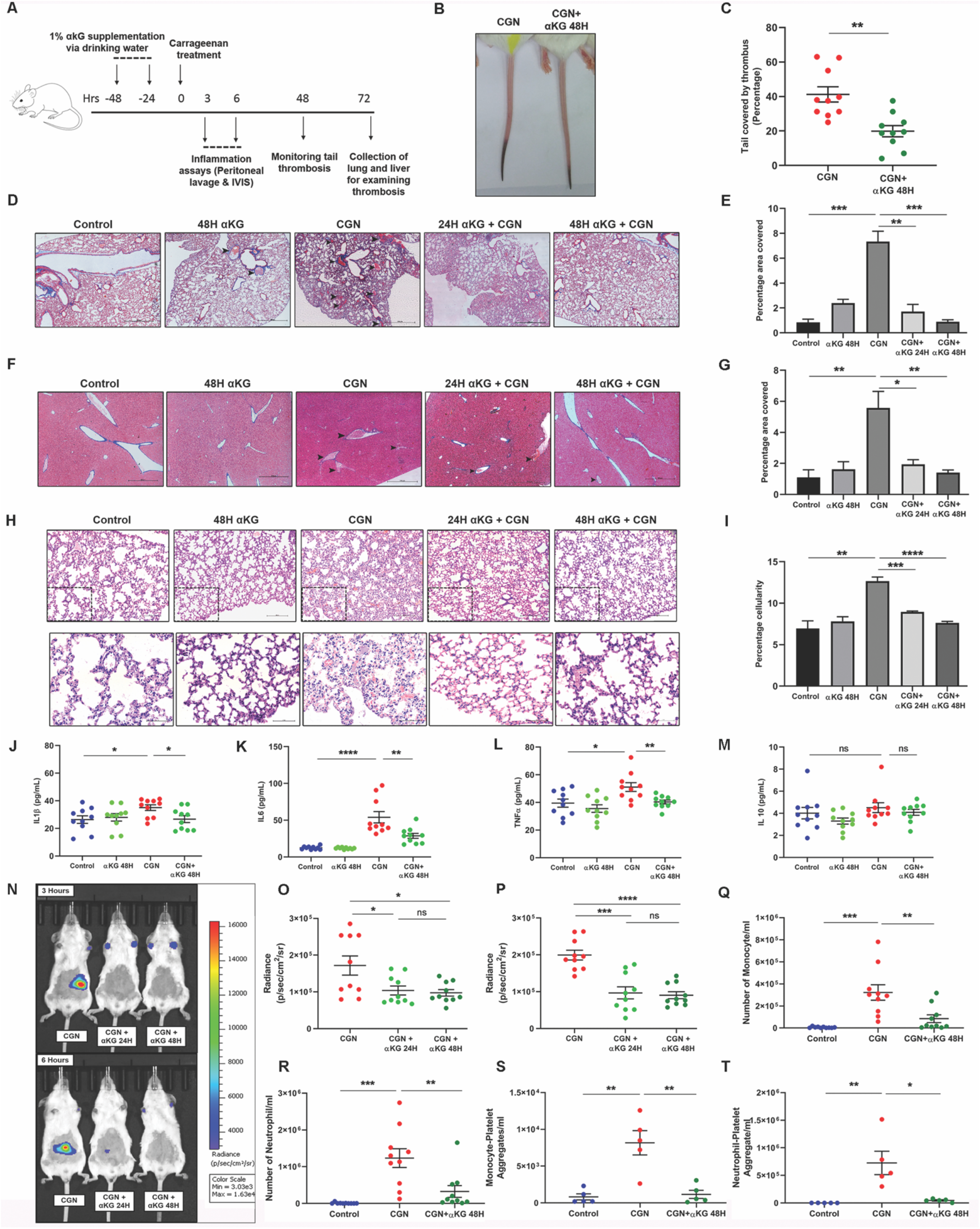
Dietary *α*KG supplementation rescues mice from carrageenan-induced thrombosis and inflammation. (**A**) Schematic showing experimental protocol. BALB/c mice were supplemented with dietary αKG before exposed to carrageenan treatment and following parameters were assessed. (**B-C**) (B) Tail thrombosis was measured, showing a representative image. (C) Percentage thrombosis. Each dot represents a mouse. (**D-G**) (D-E) Lung and (F-G) Liver section from above mice groups were used for assessing clot formation using Masson-trichrome (MT) staining. Scale bar 500 µm. Percentage covered area of clot was measured. The hematoxylin and eosin (H&E) staining data of similar observations are mentioned in Supplemental Figure 10. (**H-I**) Similarly, H&E staining of lung was used for assessing leukocyte accumulation. Score is calculated as percentage cellularity. Scale bar 500 µm and 20 µm. Showing reduced inflammation in αKG-treated mice. The MT staining data of similar observations are mentioned in Supplemental Figure 11. Figure D-I, n=8. (**J-M**) Cytokines (J) IL1β, (K) IL6, (L) TNFα And (M) IL10 were measured from mice groups. (**N-T**) To measure local inflammation αKG-supplemented mice were injected with carrageenan at peritoneum region and (N) luminescence generated from MPO-luminol interaction were measured using IVIS imaging system to assess leukocyte activation and inflammation. Radiance count from (O) 3 and (P) 6 hours post carrageenan treatment. αKG-supplemented mice displayed reduced MPO signal intensity. (Q) Monocytes, (R) Neutrophils, (S) monocytes-platelet and (T) neutrophil-platelet aggregates were measured from peritoneum lavage of above treated mice. Each dot represents a mouse. Data in above figure are Mean ± SEM from different experiments. Unpaired t-test was used to compare between the groups, \**P*<0.05, \*\**P*<0.01, \*\*\**P*<0.001, \*\*\*\**P*<0.0001 and ns=non-significant.

### Dietary αKG significantly inhibits inflammation and thrombosis alongside downmodulation of pAkt in lung of the hamsters infected with SARS-CoV-2

A recent report suggested the increased expression of pAkt1(Thr308) in human alveolar epithelial type 2 cells after SARS-CoV-2 infection (17), we also observed a similar elevated expression of pAkt1(Thr308) in human liver Huh7 cell line infected with this virus. As expected, we observed a significant reduction in pAkt1(Thr308) expression along with decreased IL6 secretion in these infected cells after octyl-αKG supplementation (Figure 6A-C), but viral load did not change significantly (Figure 6D). We then tested the rescue effect of dietary αKG (1%), administered via drinking water and oral gavage (protocol of treatment is mentioned in Figure 6E), and observed significantly reduced clot injury spots on lung in SARS-CoV-2 infected hamsters (Figure 6F). The histopathology data showed a significantly reduced intravascular clot formation (Figure 6G-H) and leukocytes accumulation in alveolar spaces (Figure 6I-J) in the lung of SARS-CoV-2 infected hamsters after αKG administration. The elevated expression of pAkt in lung of SARS-CoV-2 infected animals was significantly reduced after αKG supplementation (Figure 6K-L). Besides, an elevated expression of HIF2α in the lung of SARS-CoV-2 infected animals was significantly reduced after αKG supplementation (Supplemental Figure 13A-B). The body weight decreased significantly in infected animals during day 3 - day 6, but no rescue effect of αKG on body weight was observed (Figure 6M). As reported by others (27), we did not observe death of SARS-CoV-2 infected hamsters. We also observed a gradual increase in body weight on day 8 onwards in another group of all infected hamsters. However, a decreased viral load in the lung was observed at day 6 in infected animals supplemented with αKG (Figure 6N). Supplementation with 1% dietary αKG for 6 days to control hamsters did not alter the count of blood cells including platelet and WBCs and granulocytes (Supplemental Figure 8D-F), suggesting a safe implementation of the metabolite.

**Figure 6.**
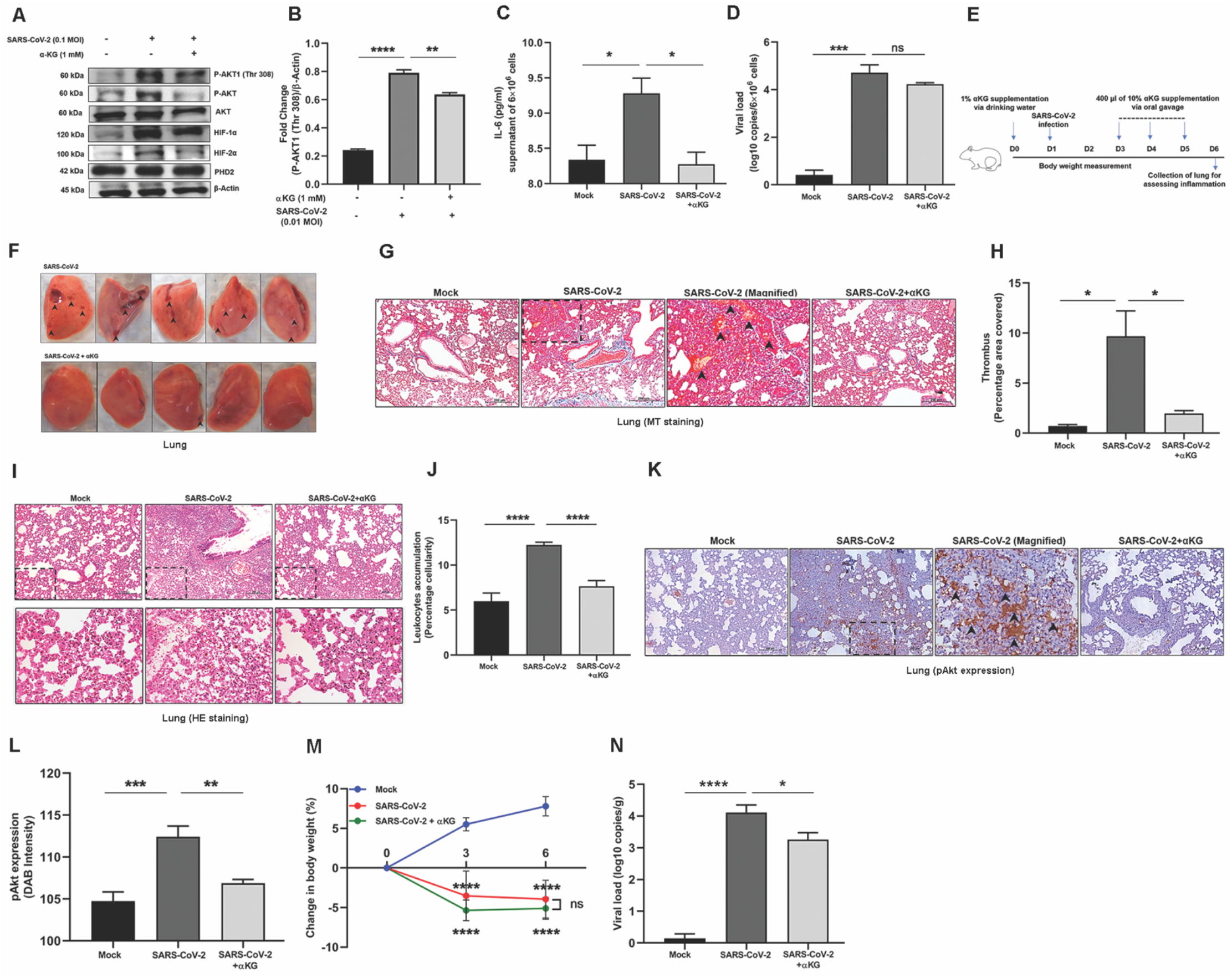
Dietary *α*KG supplementation rescues hamsters from SARS-CoV-2 induced lung thrombosis and inflammation. (**A-B**) Before performing *in vivo* study we performed *in vitro* experiment and infected human liver Huh7 cell line with 0.1 MOI of SARS-CoV-2 for 24 hrs with a prior treatment with 1mM octyl-αKG. (A) We assessed pAkt1(Thr308) expression using WB. (B) Densitometry analysis from three independent experiments shows elevated expression of pAkt1(Thr308), other densitometry data are mentioned in Supplemental Figure 14M-P. (**C**) IL6 level was measured from supernatant of SARS-CoV-2 infected Huh7 cells from above experiment. (**D**) SARS-CoV-2 genome was measured from above cell pellet using RT-PCR. (**E**) Schematic Figure shows the experimental protocol using hamsters. (**F**) Lung was isolated and imaged, showing clot injury spot (arrows) after viral infection, which was rescued to almost normal condition with αKG supplementation. (**G-H**) (G) Lung section from above mice groups were used for assessing clot formation and collagen deposition using MT staining and (H) Percentage covered area of thrombus was measured. (**I-J**) (I) H&E staining of lung was used for assessing leukocyte accumulation in lung including alveolar spaces as the marker of inflammation. (Lower panel) magnified version from upper panel images. (J) Score is calculated as percentage cellularity. Showing reduced inflammation in αKG-treated animals. (**K-L**) (K) The sections were labelled with pAkt antibody and stained with HRP-conjugated secondary antibody. (L) pAkt expression was quantified by measuring DAB staining (brown), which was increased in virus infection and decreased after αKG treatment. (G-L) Image scale bar 200 µm and number of animals used n=5 in mock, n=9 each in other groups. (**M**) Body weight of the animals were measured at day 0, 3 and 6, and calculated as Mean ± SD, n=5 in mock, n=11 each in other groups. (**N**) SARS-CoV-2 genome was measured from lung tissue (gram) of hamsters using RT-PCR, n=3 in mock, n=11 each in other groups. Data in above figure are Mean ± SEM from independent experiments. Unpaired t-test was used to compare between the groups, \**P*<0.05, \*\**P*<0.01, \*\*\**P*<0.001, \*\*\*\**P*<0.0001 and ns=non-significant.

## Discussion

Our study for the first time describes the regulatory role of PHD2-pAkt axis in platelet function. Also, we show heretofore unreported inactivating impact of α-ketoglutarate (αKG) mediated augmentation of prolyl hydroxylation activity of PHD2 on phosphorylated Akt1 (pAkt1). An earlier study has described that pAkt1 undergoes prolyl hydroxylation at Pro125 and Pro313 by PHD2 in a reaction decarboxylating αKG. Hydroxylated pAkt1(Thr308) is dephosphorylated by Von Hippel-Lindau protein (pVHL) associated protein phosphatase 2A (PP2A), leading to Akt inactivation. Study unveils this pathway as another line of post translational modification for pAkt. In VHL-deficient/suppressed setting and under hypoxic microenvironment, accumulation of pAkt is likely to promote tumour growth and its inhibition partially reverses the effect (25).

We describe here that the supplementation with αKG, an intermediate of TCA cycle, significantly suppresses pAkt1 and reduces agonist-induced platelet activation under normoxia. Upon activation by agonists like collagen (3) and ADP (30) platelets undergo Akt phosphorylation by PDK1. Akt phosphorylation is known to stimulate cell surface adhesion molecules like GPIIbIIIa and GPVI, and promote platelet aggregation and adhesion, and also secretion of granular contents (31-32). αKG supplementation significantly inhibits the above –mentioned functions of platelets by suppressing pAkt1. Contrastingly, a marked amplification in platelet activity alongside increased pAkt1 was observed after treatment with PHD2 inhibitors like DKG or DHB. This indicates a crucial involvement of PHD2 in the regulation of platelet activation. The above speculation was further supported by the αKG-mediated reduced expression of HIF1α and HIF2α, known substrates of PHD2, and DKG/DHB-driven stabilization of the same. Augmented PHD2 activity orchestrated these events was confirmed by increased hydroxylated proline alongside enhanced binding of PHD2 to pAkt in αKG-treated platelets. Besides, our study also describes an increased intracellular αKG: succinate ratio in platelets after αKG supplementation. PHD2 catalyses proline hydroxylation of its substrate by converting O2 and αKG to CO2 and succinate (33), and succinate can inhibit PHD2 by competing with αKG (34). Therefore, an elevated intracellular ratio of αKG to succinate may serve as a stimulator of PHD2 activity as suggested (28). It is notable that αKG: succinate ratio was unaltered in platelet after activation with agonist, but its elevated ratio in αKG-supplemented platelet significantly augmented PHD2 activity and in turn downmodulated pAkt1. Contrastingly, several studies have reported an increased pAkt in platelets and other cells *in vitro* (35-36) and *in vivo* (37) after succinate supplementation. Thus, indicating that intracellular ratio of these two metabolites serves as a switch for the PHD2 activity. We show that αKG supplementation significantly decreased pAkt in platelet to inhibit aggregation, thrombus formation and secretion of granular contents including inflammatory mediator such as sphingosin-1 phosphate (S1P) *in vitro*. S1P is one of the connecting between platelet activation and systemic inflammation as it can activate monocytes. Interestingly we observed that αKG could also suppress S1P-mediated activation of monocytes in a pAkt1 dependent manner. When LPS was used as an activator to simulate a thrombo-inflammatory condition, αKG could deter the secretion of pro-inflammatory cytokines from monocytes. Importantly, our data showed no significant inactivation of pAkt1 by αKG in PHD2-deficient monocytic cell line, thus confirming that αKG imparts its effects primary through PHD2 in this context. We could also confirm that the outcomes of αKG usage were independent of HIF1α. Overall, our data upholds PHD2 as a potential target to abrogate Akt signalling. Studies have extensively used antagonists/inhibitors to target pathological signalling of Akt or PI3K-Akt to inhibit thrombosis and inflammation (8, 4, 23-24).

Our study highlights the implementation of dietary αKG to mice as one of the potential treatments to reduce the platelet aggregation and inflammatory response of monocyte by downmodulating pAkt1 without altering the count of these cell types. Therefore, it suggests a safe administration of this metabolite. In a recent work, we have described a safe rescue effect of this metabolite in mice from hypoxia-induced inflammation by downmodulating HIFα (29). αKG has been used extensively for *in vivo* experimental therapies for manipulating multiple cellular processes related to organ development and viability of organisms (38-39), restriction of tumor growth and extending survival (40), and preventing obesity (41).

Another interesting part of our study describes the αKG-mediated rescue of clot formation and leukocyte accumulation alongside a reduction in cytokine secretion by these cells in lung and other organs in mice exposed to a thrombosis-inducing agent like carrageenan. Specifically, our study also reports a significant rescue effect of αKG on inflamed lung in SARS-CoV-2 infected hamsters. A significant reduction in intravascular clot formation and accumulation of leukocytes including macrophages and neutrophils in alveolar spaces of the lung of infected hamsters suggests a potential rescue effect of αKG. Thus, indicating that αKG usage can decelerate inflammation induced lung tissue damage reported in severe cases of SARS-CoV-2 infection and may eventually deter development of acute respiratory distress syndrome (42-43). However, this speculation needs further experimental evidences. Besides, αKG treatment also decreased viral load in the lung of the infected animals alongside diminished expression of pAkt. Although the exact role Akt in replication of SARS-CoV-2 remains to be delineated as reported for other viruses (44). But its noteworthy that a recent study has described an elevated pAkt1(Thr308) in cells infected with SARS-CoV-2 (17). Our *in vitro* data also show an elevated pAkt1(Thr308) alongside increased IL6 secretion by SARS-CoV-2 infected Huh7 cell line, which was further inhibited by αKG administration. Thus, suggesting that the augmentation of PHD2 activity by αKG would be a potential therapeutic strategy to inhibit pAkt-mediated anomalies like inflammation and thrombosis in host and also propagation of SARS-CoV-2.

Therefore, our data together suggest a novel role of PHD2-pAkt axis in the regulation of platelet and leukocyte functions. Supplementation with αKG significantly increases the hydroxylase activity of PHD2 and therefore reduces phosphorylation of Akt and in turn supresses thrombotic and inflammatory functions of platelets and leukocytes respectively, depicted in schematic Figure 7. Thus, suggesting a safe implementation of dietary αKG in prevention of Akt-driven thrombosis and inflammation in various disease conditions including inflamed lung in COVID-19. Study also highlights αKG-PHD2-pAkt axis as a potential target for better pulmonary management in these diseases.

**Figure 7.**
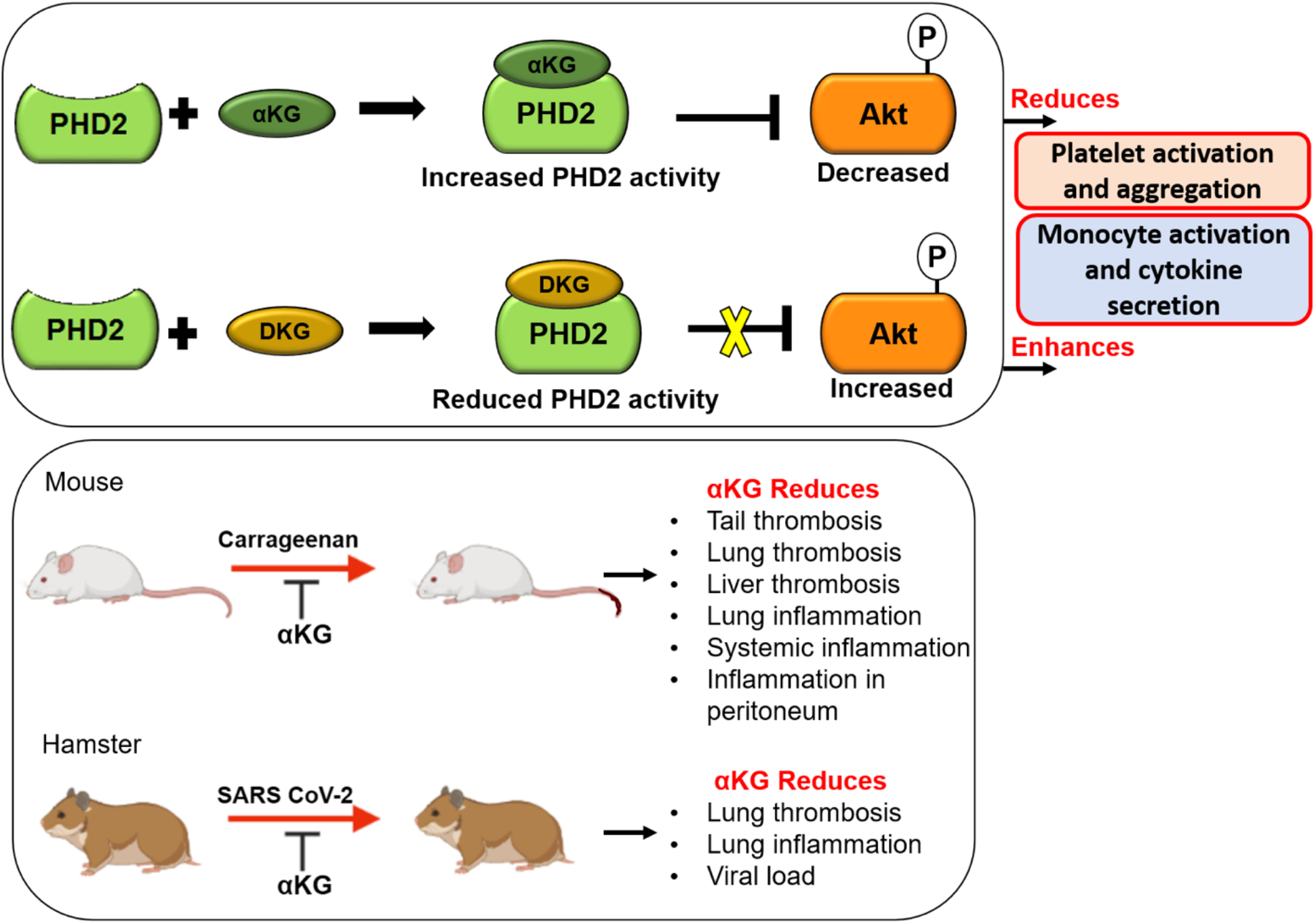
*α*KG mediated suppression of platelet and monocyte functions. Schematic depicts that under normoxic environment supplementation with αKG (known cofactor of PHD2) increases PHD2 activity by elevating intracellular αKG: succinate ratio. Elevated PHD2 activity degrades pAkt1 and reduces platelet activation and aggregation. A similar mechanism also suppresses inflammatory function of monocyte. Thus, suggesting the involvement of αKG-PHD2-Akt1 axis in the regulation of events like thrombosis and inflammation. Importantly, dietary αKG supplementation significantly rescues mice from carrageenan-induced clot formation and leukocyte accumulation and inflammation in various organs including lung. Importantly, dietary αKG rescues significantly hamsters from SARS-CoV-2 induced clot formation and leukocyte accumulation in the lung, alongside downmodulation of pAkt.

## Materials and Methods

### Platelet and monocyte isolation

Whole blood (16 ml) was collected form healthy individuals in sodium citrate or ACD anticoagulant. Platelets and monocytes were isolated from whole blood and used for *in vitro* experiments. 16 ml of whole blood was collected from healthy volunteers by venepuncture in vacutainers containing anti-coagulant sodium citrate or acid-citrate dextrose (ACD). Platelet rich plasma (PRP) was separated by centrifugation at 44 g for 15 min. Sodium citrate containing PRP was used for aggregation and activation studies. PRP in ACD was used for isolating washed platelets for further studies as described in our previous work (45).

Peripheral blood mononuclear cells (PBMCs) were isolated using Ficoll Hypaque (GE Healthcare, Freiburg, Germany) density gradient centrifugation as mentioned in our earlier work (46). PBMCs were washed twice with PBS, pH 7.4 and seeded in cell culture-treated plates (Corning, NY, USA) in RPMI-1640 medium (Sigma Aldrich, USA) supplemented with 10% (v/v) fetal bovine serum (Gibco Invitrogen, San Diego, CA), 100 U/mL penicillin and 100 µg/mL streptomycin for 2 hrs at 37°C in a humidified atmosphere with 5% CO^2^, to allow monocytes to adhere to the plate. After 2 hrs, supernatant containing non-adherent cells were removed and adhered monocytes were used for further treatments mentioned hereafter.

### Platelet activation and aggregation assays

Human PRP diluted (1:1) in Tyrod’s buffer pH 7.2 was used for following assays. Diluted PRP was pre-treated with octyl α-ketoglutarate (Sigma Aldrich, USA) or inhibitors to PHD2 such as dimethyl ketoglutarate (DKG, Sigma Aldrich) or ethyl-3-4-dihydroxybenzoic acid (DHB, TCI America, Portland) and incubated with collagen (10 µg/ml) or ADP (2 µM, both from the Bio/Data Corporation, USA) 10 minutes. Platelets were labelled with P-selectin (PE Cy5), PAC-1 (FITC) and Annexin-V (FITC) antibodies (BD bioscience) for 15 min at 37°C and fixed in 1% paraformaldehyde. 20,000 events were acquired using flow cytometry (BD FACS Verse). The acquired data was analysed using the Flowjo software (Tree Star, USA), as mentioned (45).

Platelet aggregation was performed using PAP8 aggregometer (Bio/Data Corporation, USA). PRP was pre-treated with αKG or DKG and incubated with collagen (20 µg/ml) or ADP (5 µM) and aggregation percentage was measured.

### Platelet thrombus formation assay

Platelet thrombus formation assay was performed by perfusing whole blood (collected in citrate-anticoagulant from healthy individuals) over the petri plate immobilized with collagen. Whole blood was preincubated for 5 min with either αKG or DKG or both before perfusion on collagen coated surface. A syringe pump (Harvard Apparatus Inc., USA) was connected to the outlet port that drew blood through the chamber at arterial shear stress of 25 dyne/cm^2^. The flow chamber was mounted onto a Nikon Eclipse Ti-E inverted stage microscope (Nikon, Japan) equipped with a high-speed digital camera. Movies were recorded at magnification 40X and analysed using NIS-Elements version 4.2 software as mentioned in our previous study (47).

### Platelet microparticle measurement

PRP was pre-treated with/without αKG and incubated with collagen for 10 min. Platelet-free plasma was obtained by 2 sequential centrifugations: PRP at 1500 g for 7 min followed by platelet-poor plasma (PPP) at 1500 g for 15 min. Platelet-derived microparticles (MPs) were measured using flow cytometry after labelling with anti-CD41 PE antibody as mentioned (45).

### Monocyte activation

Primary monocytes and monocytic cell line were pre-treated with 1mM octyl α-ketoglutarate (Sigma Aldrich, USA) for 2 hrs and 4 hrs respectively followed by replacement with fresh media. Cells were treated with either S1P (1 µM) or LPS (500 ng/ml). Treated cell supernatant was used for assessing cytokines using the cytometric bead array (CBA). Protein lysate prepared from cell pellet was used for western blotting of signalling molecules.

### CBA for quantifying cytokines

Cytokines such as TNF-α, IL-1β, IL-6 and IL-10 were measured from human primary monocytes culture supernatant or mice plasma of different treatments as mentioned in results using CBA and analysed by FCAP array software (BD Biosciences, San Jose, CA,USA).

### Generation of HIF-1*α* depleted monocyte cell line

HIF-1α protein expression was depleted in human U937 cell line (ATCC, USA) using shRNA targeting HIF-1α (TCRN0000010819, Sigma Aldrich, USA) using a liposome mediated delivery (Life Technologies, Thermo Fisher Scientific, USA).

### Generation of PHD2 depleted monocyte cell line

PHD2 was depleted in human U937 cell line using shRNA targeting *EGLN1* as mentioned our publication (29).

### Dietary *α*KG supplementation to mice and hamsters

Male BALB/c mice aged 5-7 weeks or Syrian golden hamsters of 8 weeks were supplemented with 1% of dietary α-ketoglutarate (SRL, Mumbai, India) in drinking water for 24 or 48 hrs (to mice), or for 6 days (to hamsters) as mentioned in schematic Figure 4, 5 and 6. The blood cell counts and other assays were performed.

### Mice platelet aggregation

PRP was collected from control and αKG-treated mice and diluted with PBS (1:1 vol) and processed for centrifugation at 90 g using brake-free deceleration on a swinging bucket rotor for 10 min. Platelets counts were adjusted to 2.5×10^8^/ml and platelet aggregation was performed using PAP8 aggregometer. Collagen (7.5 µg/ml) or ADP (5 µM) was used as aggregation agonist.

### Carrageenan treatment to mice and measuring thrombosis and cellularity scores

BALB/c mice were used to develop carrageenan-induced thrombosis model (26). Mice were injected with 100 µl of 10 mg/ml κ-carrageenan (Sigma Aldrich, USA) prepared in normal saline in intraperitoneal cavity. αKG was supplemented via drinking water to these mice. After 48 hrs of carrageenan treatment, length of thrombus covered tail was measured and percentage tail thrombosis was calculated by length of thrombus covered tail/total length of tail×100.

Thrombosis score was measured in lung and liver of above mice. Lungs and liver samples were fixed in 4% formalin and paraffin embedded. 2.5 µm thick sections were prepared and stained with Haematoxylin and Eosin (H&E) and Masson’s trichrome (MT). Slides were observed under Nikon Eclipse Ti-E inverted stage microscope (Nikon, Japan) and images were acquired at 20 X to observe thrombi and at 40 X to observe leukocyte accumulation. Thrombosis scoring was calculated using ImageJ software. Thrombi were selected using freehand selection tool in MT stained slides. Percentage area covered was calculated as percentage of freehand selected area covering the total area. Similarly, leukocyte accumulation was assessed as a marker of inflammation in the above lung section using percentage cellularity. Cellularity score was calculated using ImageJ software. Images were converted to RGB stack and from that all nuclei were selected based on the intensity of color and size from H&E stained slides. Percentage area covered by nucleated cells was calculated by measuring nuclear area as a percentage of the total tissue section area.

Carrageenan-induced peritoneal inflammation was measured in mice using a modified protocol as described (48). BALB/c mice received carrageenan (10 mg/ml) or saline intraperitoneally. At 3hrs and 6hrs, the animals were anaesthetized and peritoneal exudates were harvested in 3 ml of PBS. Different immune cell populations in peritoneal lavage were analysed and counts were determined by flow cytometry using CD45.2, CD11b, CD11c, Ly6G, Ly6C, and CD41 (46, 49). In another set of a similar experiment, BALB/c mice were injected with carrageenan. At 3 and 6 hrs, peritoneal inflammation was visualised using bioluminescence based imaging of MPO activity by injecting luminol (i.p. 20 mg/100 g body weight) (Sigma Aldrich) 6 min prior to imaging using an in-vivo imaging system (IVIS; Perkin Elmer, Waltham, MA, USA) as mentioned in our work (46).

### SARS-CoV-2 infection to Syrian golden hamsters

Male golden hamsters of 8 weeks old were given infection of SARS-CoV-2 (isolate USA-WA-1/2020 from World Reference Center for Emerging Viruses and Arboviruses, from UTMB, Texas, USA), via nasal route inoculation using 1 × 10^6^ plaque-forming units (PFU) as mentioned (27). 1% dietary αKG was administered via drinking water and 400µl of 10% αKG was given through oral gavage on day 3 through day 5. During this phase they were symptomatic and were not drinking sufficient (male hamster of 100 gm B wt. drinks normally 5 ml per day) water. The schematic protocol of infection and therapy is mentioned in Figure 6. At day-6, animals were sacrificed and lungs and liver samples were harvested, fixed in 4% formalin, paraffin embedded and processed for H&E and MT staining. The thrombosis and inflammation scores were measured as mentioned above. The lung sections were used for immunohistochemistry staining for pAkt (Cell Signalling Tech, USA) Body weight was recorded on alternate days. The lung sections were used for measuring viral genome using RT-PCR.

### SARS-CoV-2 infection to cell line

Human liver cell line Huh7 (ATCC, USA) was seeded (6×10^4^ cells/well) and pre-treated with 1mM octyl αKG for 2 hrs and infected with 0.1 MOI of SARS-CoV-2 for 24 hrs in BSL3 facility. The cells pellet was lysed and fixed for using estimation of pAkt1 using western blotting.

### RT-PCR detection of viral genome

Lung tissue sample from hamsters was homogenized in Trizol reagent (MRC, UK) using a hand-held tissue-homogenizer and the total RNA extracted as per manufacturer’s protocol. 1 µg total RNA was reverse-transcribed using Superscript-III reverse-transcriptase (Invitrogen, USA) as per manufacturer’s protocol, using random hexamers (Sigma Aldrich, USA). The cDNA was diluted in nuclease-free water (Promega, USA) and used for real-time PCR with either SARS-CoV-2 or GAPDH specific primers, using 2x SYBR-green mix (Takara Bio, Clontech, USA) in an Applied Biosystems® QuantStudio(tm) 6 Flex Real-Time PCR System. The oligonucleotides used were SARS-F (5’-CAATGGTTTAACAGGCACAGG-3’) and SARS-R (5’-CTCAAGTGTCTGTGGATCACG-3’) for SARS-CoV-2, and G3PDH-F (5’-GACATCAAGAAGGTGGTGAAGCA-3’) and G3PDH-R (5’-CATCAAAGGTGGAAGAGTGGGA-3’). The Ct value corresponding the viral RNA was normalised to that of G3PDH transcript. The relative level of SARS-CoV-2 RNA in mock-infected samples was arbitrarily taken as 1 and that of infected samples expressed as fold-enrichment (FE). The FE value for each infected sample was transformed to their logarithmic value to the base of 10 and plotted.

The viral genome was measured in Huh7 cell line treated with SARS-CoV-2 using RT-PCR as mentioned above. The Ct value corresponding the viral RNA was normalised to that of RNAse P (RP) transcript. The oligonucleotides used were RP-F (5’-AGATTTGGACCTGCGAGCG-3’) and RP-R (5’-GAGCGGCTGTCTCCACAAGT-3’).

### Immunoprecipitation

PHD2 was immunoprecipitated from platelet lysate using protein G sepharose beads and anti-PHD2 antibody (Cell Signalling Tech, USA). Similarly, phosphorylated Akt (pAkt) was immunoprecipitated using protein A sepharose beads and anti-pAkt antibody (Cell Signalling Tech, USA). Briefly, washed platelets isolated from whole blood of healthy volunteer and lysed in lysis buffer (25 mM Tris, 150 mM sodium chloride, 1 mM EDTA, 1% NP-40, 50 mM sodium fluoride and 3% Glycerol) with protease and phosphatase inhibitor. Lysate was pre-cleared with protein A or G sepharose beads and then added to antibody coated beds and incubated overnight.

Beads were removed, washed with lysis buffer and collected protein sample was processed for western blotting.

### Western blotting

The whole cell (platelets or primary monocyte or monocytic cell line) lysate was prepared using RIPA lysis buffer and protease-phosphatase inhibitor (Thermo Scientific Life Tech, USA). SDS-PAGE gel was followed by immunoblotting using primary antibodies against pAkt, Akt, pAkt1(Thr 308), Akt1, HIF-1α, HIF-2α, β-Actin (Cell Signalling, USA) and α-tubulin (Thermo Fisher Scientific, USA) as described in detail in our previous work (45). The detailed information of antibodies is mentioned in Supplemental Table 1.

### Estimation of metabolites

Steady-state level of α-ketoglutarate, Lactate, Fumarate, Pyruvate and Succinate was estimated in plasma, PBMC-granulocytes (10^5^) and platelets (10^5^) of mice or human samples from different treatments as per manufacture protocol (Sigma Aldrich, USA catalog no. MAK054, MAK064, MAK060, MAK071, MAK335 respectively).

### Sphingosine-1-Phosphate measurement

αKG treated and control mice platelets were stimulated with collagen (10 µg/ml) *in vitro* and supernatant was collected and used to estimate Sphingosine-1-phosphate (S1P) level as per manufacturer’s protocol [Cloud clone corp. (CEG031Ge)]. Similarly, human platelets were treated with αKG and collagen (5 µg/ml) and supernatant was used to estimate S1P level.

### Statistical Analysis

Data from at least three experiments are presented as Mean ± SEM (Standard Error of the Mean). Statistical differences among experimental sets were analysed by Unpaired t test. Graph Pad Prism version 8.0 software was used for data analysis and *P*-values **<**0.05 were considered statistically significant.

## Supplemental Materials

Supplemental Figure 1: Measuring: Akt1 phosphorylation (pAkt1) in platelet induced by ADP; agonist induced pAkt1 in platelet in presence of PHD2 inhibitor DHB; and pPI3K inn presence of αKG.

Supplemental Figure 2: IgG control for immunoprecipitation of PHD2 and P-Akt from platelet lysate.

Supplemental Figure 3: Microparticle release from platelets after αKG treatment.

Supplemental Figure 4: ADP induced platelet aggregation in presence of αKG, DKG and DHB.

Supplemental Figure 5: Effect of αKG on LPS induced HIF-2α and P-Akt expression in monocytes.

Supplemental Figure 6: Generation of HIF-1α depleted cells.

Supplemental Figure 7: Measuring intracellular level of fumarate and pyruvate in activated platelets after αKG supplementation.

Supplemental Figure 8: Mice and hamster blood counts after dietary αKG supplementation.

Supplemental Figure 9: *In vitro* platelet aggregation in presence of octyl αKG, platelets were collected from normal mice; and platelet aggregation collected from αKG treated mice.

Supplemental Figure 10: H&E staining of mice lung and liver showing reduction of carrageenan induced thrombosis by dietary α-KG supplementation.

Supplemental Figure 11: Masson’s trichrome (MT) staining of mice lung showing reduction of leukocyte infiltration in carrageenan-treated mice after dietary αKG supplementation.

Supplemental Figure 12: Gating strategy for immune cells from mice peritoneal lavage fluid. Supplemental Figure 13: HIF-2α expression in lung tissue of hamsters infected with SARS-CoV-2.

Supplemental Figure 14: Densitometry of western blots. Supplemental

Supplemental Table 1: List of antibodies used in the study.

## Study approval

### Human samples

Ethics approval was obtained from the Institutional Ethics Committee (IEC) for human research of Regional Centre for Biotechnology (RCB; ref no. RCB-IEC-H-08) to recruit healthy volunteers. Volunteers were recruited on the basis of inclusion criteria: 1) healthy, 2) not taking any anti-platelet or anti-inflammatory drugs, 3) no major illness or chronic disease, and 4) no microbial infections within a month of recruitment. A written informed consent was received from all participants.

### Animal study

Ethics approval was obtained from the Institutional Animal Ethics Committee (IAEC) of RCB (ref. no. RCB/ IAEC/2020/077) and mice experiments were conducted within the guidelines of IAEC in the Small Animal Facility (SAF) of our institute. Ethics approval was obtained from the IAEC (ref. no. RCB/IAEC/2020/069) and Institutional Biosafety Committee (IBSC; ref. no. RCB/IBSC/20-21/221) of RCB and hamster experiments were conducted within the guidelines of IAEC in the BSL3 facility of SAF of our institute.

## Authors contributions

NMS designed and performed all experiments, analysed data and wrote the manuscript. SA designed and performed mice experiment for immunophenotyping and histochemistry and analysed related data. SK and Sankar B have designed and performed covid-19 infection studies, and Sankar B has supervised the infection experiments. Sulagna B designed and performed metabolite estimation and analysed data, and edited the manuscript. JTP provided crucial conceptual inputs. PG designed and supervised the study, conceptualized the approach, designed the experiments, analyzed the data, and wrote the manuscript. All authors read, edited and approved the final manuscript.

## Acknowledgements

Authors acknowledge generous help of Prof. Sudhanshu Vrati of Regional Centre for Biotechnology, Faridabad, India for sharing SARS-CoV-2 virus strain. Authors also acknowledge the funding by grants: BT/PR22881 and BT/PR22985 from the Department of Biotechnology (DBT), Govt. of India; and CRG/000092 from the Science and Engineering Research Board, Govt. of India to PG.

