## Supplementary material for "α-ketoglutarate augments prolyl hydroxylase-2 mediated inactivation of phosphorylated-Akt to inhibit induced-thrombosis and inflammation": Supplementary Information-bioRxiv-2.pdf

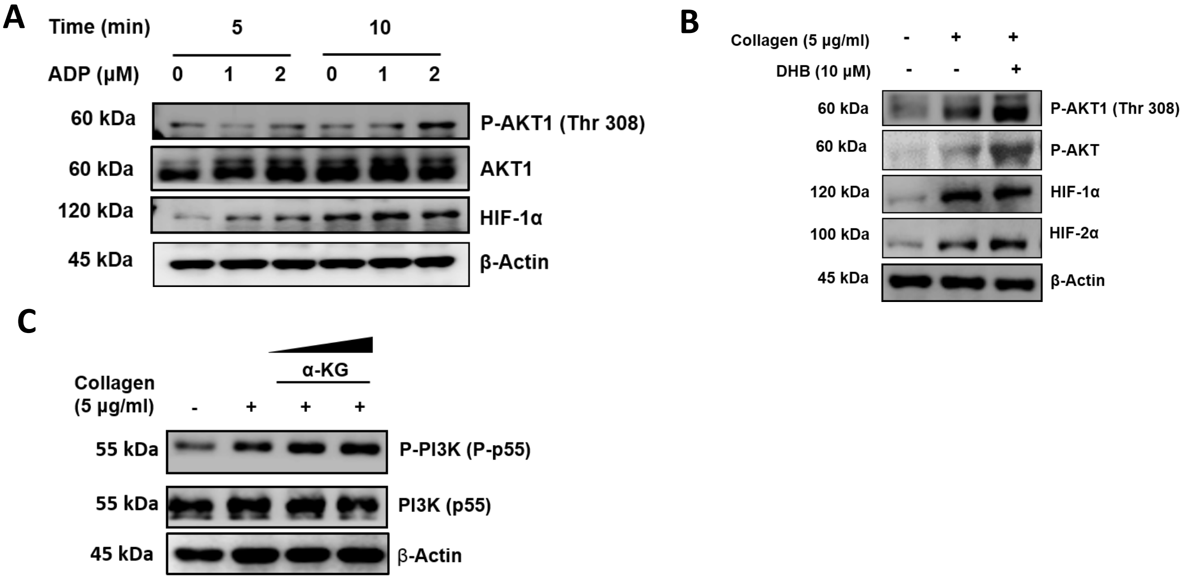


**S[illegible]ementary Figure S1. Platelet activation to A[illegible] and DHB. (A)** Platelet-rich plasma (PRP) was incubated with ADP in a time- and concentration-dependent manner and expression of phosphorylated Akt1(Thr308), Akt1, and HIF-1α were measured in platelet pellet using WB assay. Densitometry analysis shows elevated expression of P-Akt1 and HIF-1α after ADP stimulation, densitometry data are mentioned in *Online Supplementary Figure* S14Q-R. **(B)** PRP treated with collagen or DHB or both at mentioned concentration and platelet pellet was processed for WB. DHB found to increase P-Akt1(Thr308), P-Akt, HIF-1α and HIF-2α expression in platelets stimulated by collagen, densitometry data are mentioned in *Online Supplementary Figure* S14S-V. **(C)** PRP treated with collagen or αKG or both and platelet pellet was processed for WB of PI3K (p55). Densitometry data shows αKG do not affect collagen induced PI3K phosphorylation in platelets, densitometry data are mentioned in *Online Supplementary Figure* S14W.

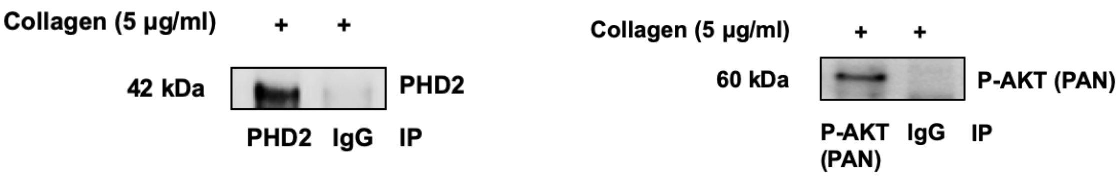


**Supplementary Figure S2. Immunoprecipitation of PHD2 and P-Akt from platelet lysate.** Washed platelets were treated with collagen (5 µg/ml) and collected platelet pellet was lysed and immunoprecipitation was performed using protein A and protein G sepharose beads for PHD2 and P-Akt (Pan). Western blot images show immunoprecipitation of **(A)** PHD2 and **(B)** P-Akt (Pan) with their respective IgG controls.

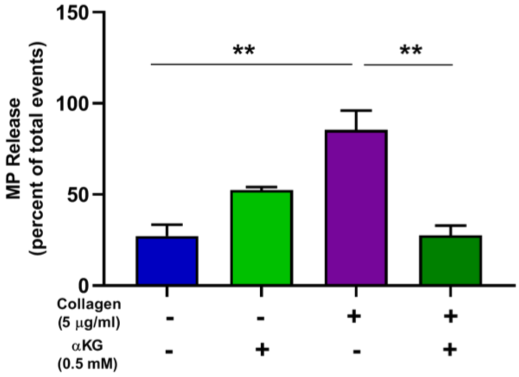


**Supplementary Figure S3. Microparticles (MP) release from platelets after αKG treatment.** Washed platelets were treated with collagen (5 µg/ml) or αKG (1 mM) or both, and collected supernatant processed for CD41a labelled microparticle analyses using flow cytometry. Bar diagram represents reduced microparticles release from α-KG treated platelets. Data are mean ± SEM from 3 independent experiments. Unpaired t-test was used to compare between the groups, ***P*<0.01.

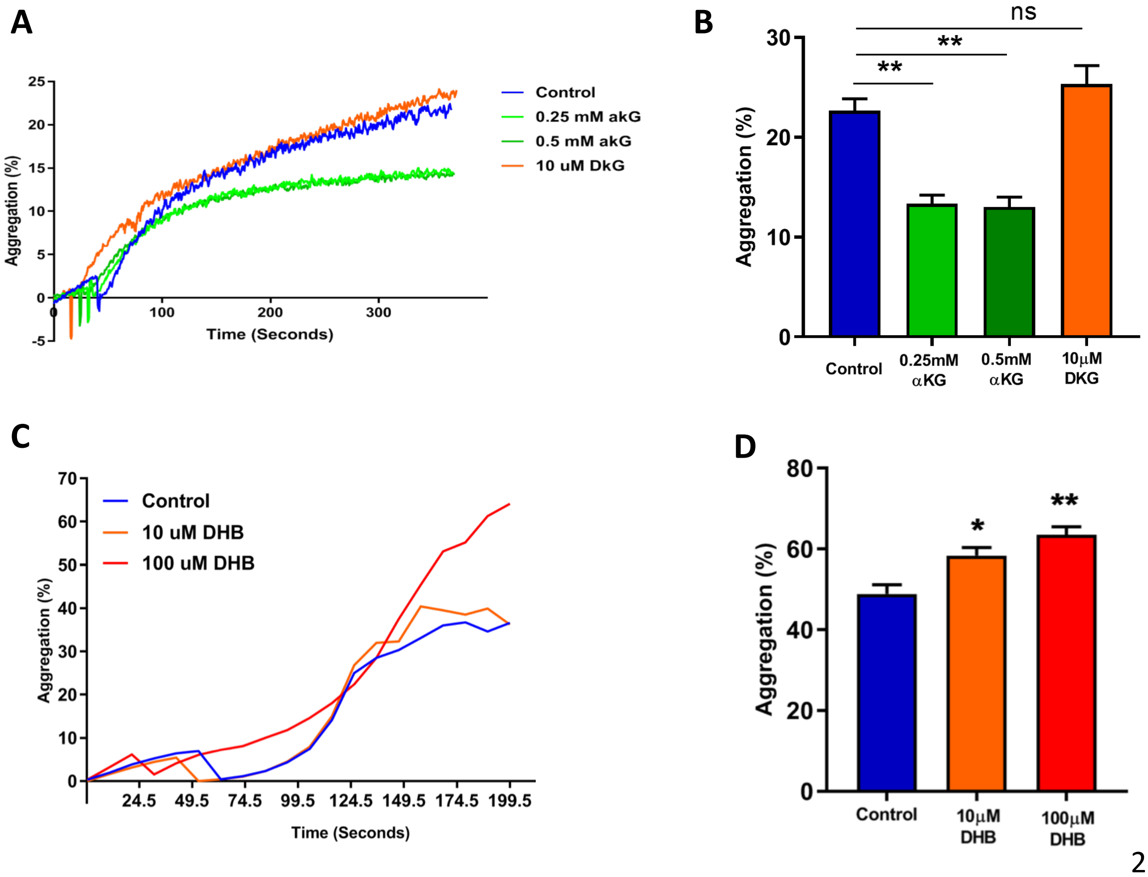

**Supplementary Figure S4. ADP induced platelet aggregation in presence of αKG, DKG and DHB. (A)** Aggregation response of platelets with ADP as aggregation stimulus was reduced upon treatment with αKG whereas DKG increases it. **(B)** Percentage platelet aggregation with ADP. **(C)** Aggregation response of human platelets with collagen as aggregation stimulus was increased with DHB treatment. **(D)** Percentage platelet aggregation with collagen. Data are mean ± SEM from 3 independent experiments. Unpaired t-test was used to compare between the groups, *$P$<0.05, **$P$<0.01, ns=non-significant.

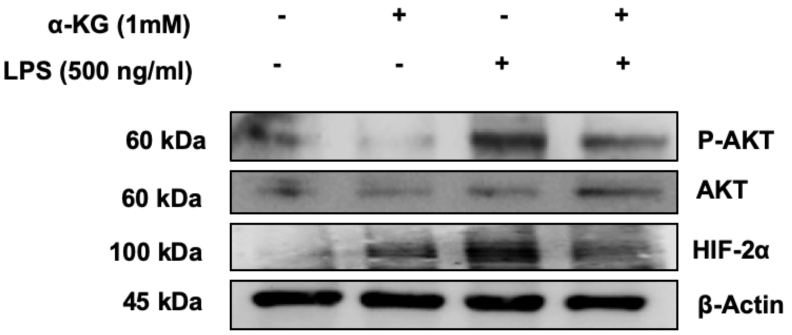


**Supplementary Figure S5. Effect of αKG on LPS induced HIF-2α and P-Akt expression.** Expression of P-Akt and HIF-2α are reduced in primary monocytes pre-treated with α-KG. Densitometry data in *Online Supplementary Figure* S14X-Y.

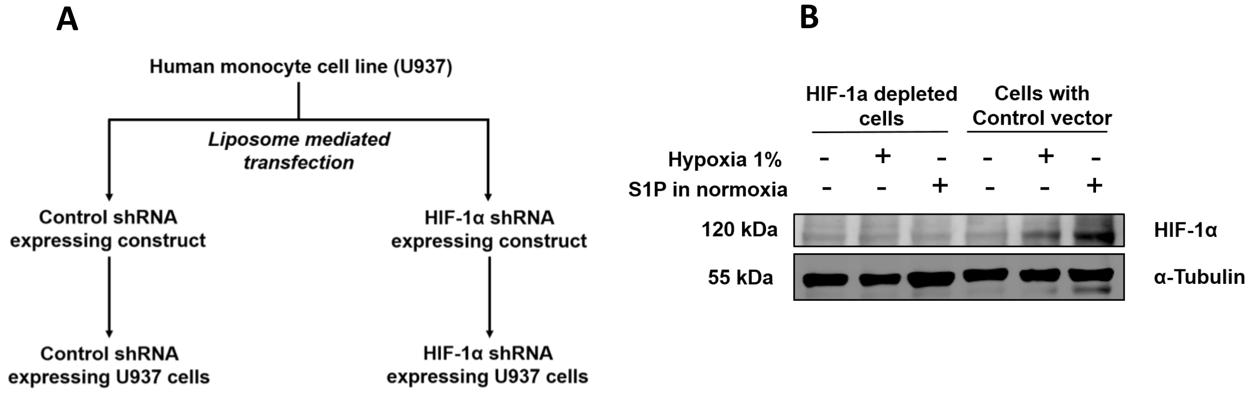

**Supplementary Figure S6. Generation of HIF-1α depleted cells. (A)** Schematic protocol for depletion of endogenous HIF-1α using shRNA in U937 cell line, using liposome-mediated delivery of construct. **(B)** HIF-1α protein expression in pLKO control shRNA and HIF-1α targeting constructs transfected cells with 24 hour 1% hypoxia or 1 μM S1P stimulation.

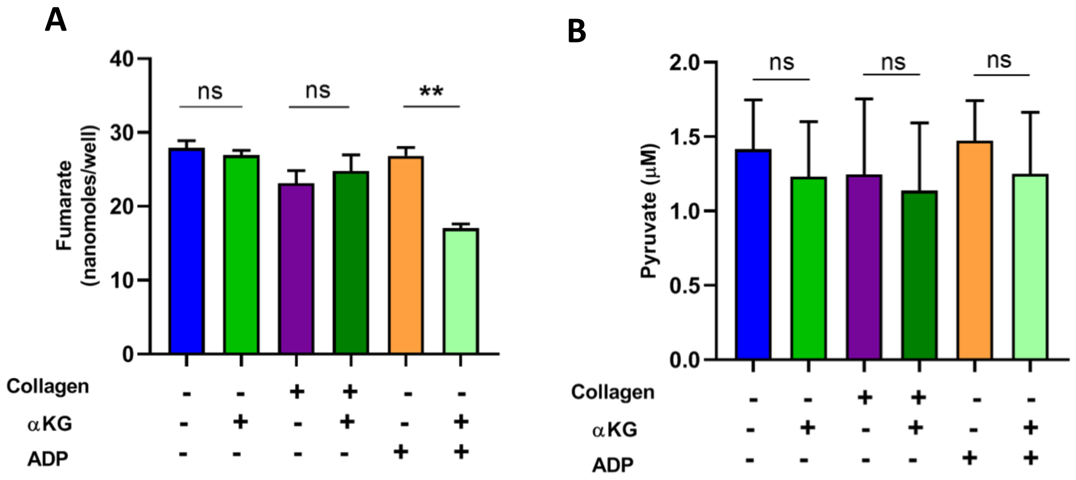


**Supplementary Figure S7. Intracellular level of fumarate and pyruvate in activated platelets after αKG supplementation.** Steady state level of **(A)** fumarate and **(B)** pyruvate were estimated from platelets lysate, pre-treated with α-KG and activated with collagen or ADP, as mentioned in Figure 3. Data are mean ± SEM from 3 independent experiments. Unpaired t-test was used to compare between the groups, ***P*<0.01, ns=non-significant.

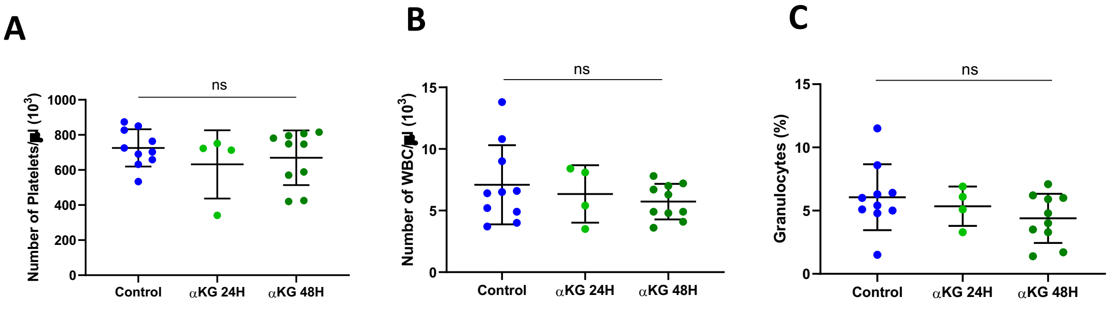

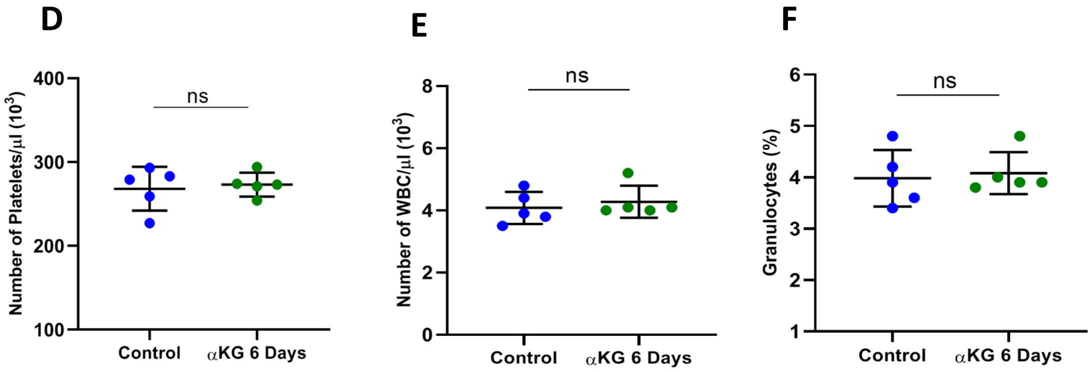


**Supplementary Figure S8. Mice and Hamster blood counts.** Count of **(A)** platelets, **(B)** WBC and **(C)** granulocytes determined from mice supplemented with dietary αKG for 24 and 48 hours, as mentioned in Figure 4A. Count of **(D)** platelets, **(E)** WBC and **(F)** granulocytes determined from hamsters supplemented with dietary αKG for 6 days (as mentioned in Figure 6), using Nihon Kohden's CelltacF haematology analyser. Data are mean ± SEM. Unpaired t-test was used to compare between the groups, ns=non-significant.

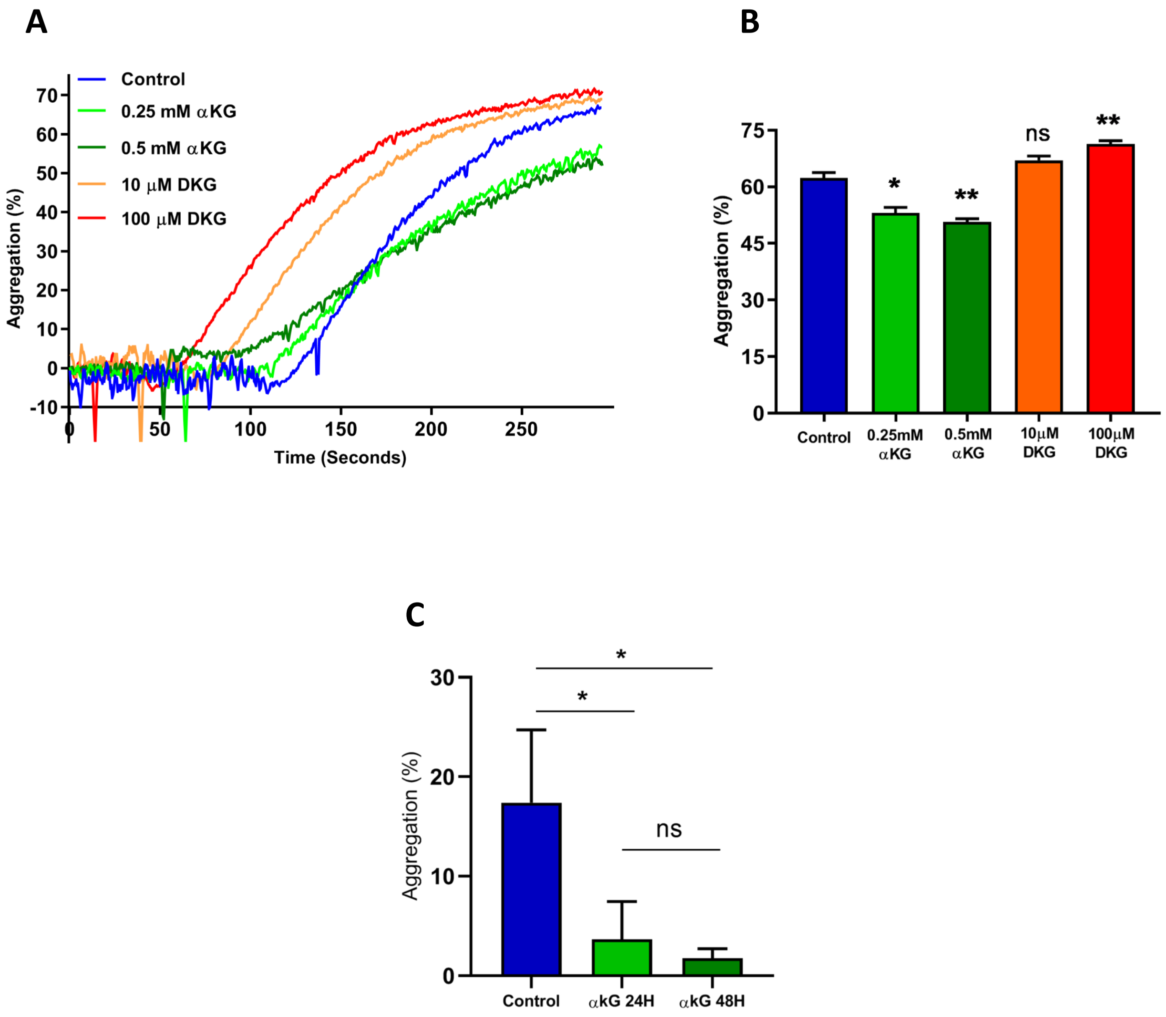

**Supplementary Figure S9. Aggregation response of mice platelets with *in vitro* and *in vivo* treatment of αKG. (A)** Aggregation response of mouse PRP to collagen with/without αKG or DKG treatment *in vitro*. **(B)** Percentage platelet aggregation. **(C)** Platelets from mice supplemented with dietary αKG for 24 and 48 hours were stimulated with ADP and aggregation response was quantified as percentage aggregation, as mentioned in Figure 4B. Data are mean ± SEM from 3 independent experiments. Unpaired t-test was used to compare between the groups, **P*<0.05, ***P*<0.01, ns=non-significant.

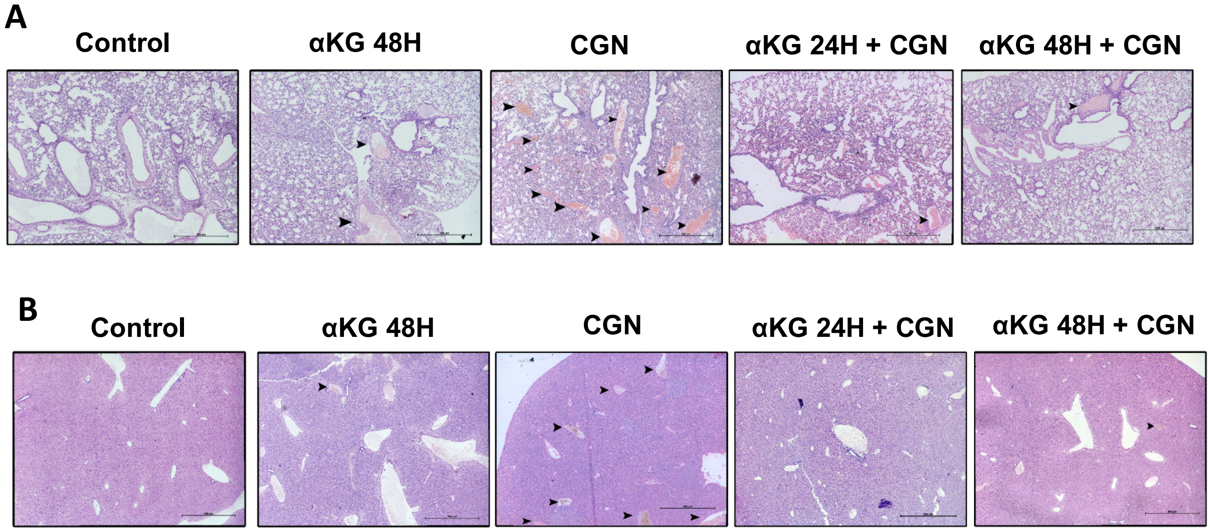


**Supplementary Figure S10. Dietary αKG supplementation reduces lung and liver thrombosis in mice.** H&E staining of mice **(A)** lung and **(B)** liver showing reduction of carrageenan induced thrombosis by dietary α-KG supplementation. Scale bar 500 µm. Similar MT staining data are mentioned in Figure 5D. Treatment details are mentioned in Figure 5A.

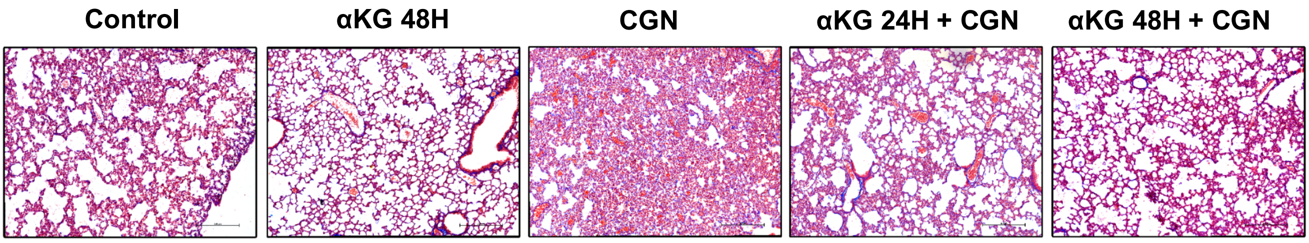


**Supplementary Figure S11. Dietary αKG supplementation reduces inflammation in mice lung.** Masson's trichrome (MT) staining of mice lung showing reduction of leukocyte infiltration, as the marker of inflammation, after dietary αKG supplementation in carrageenan-treated mice. Scale bar 500 µm. Similar H&E staining data are mentioned in Figure 5H.

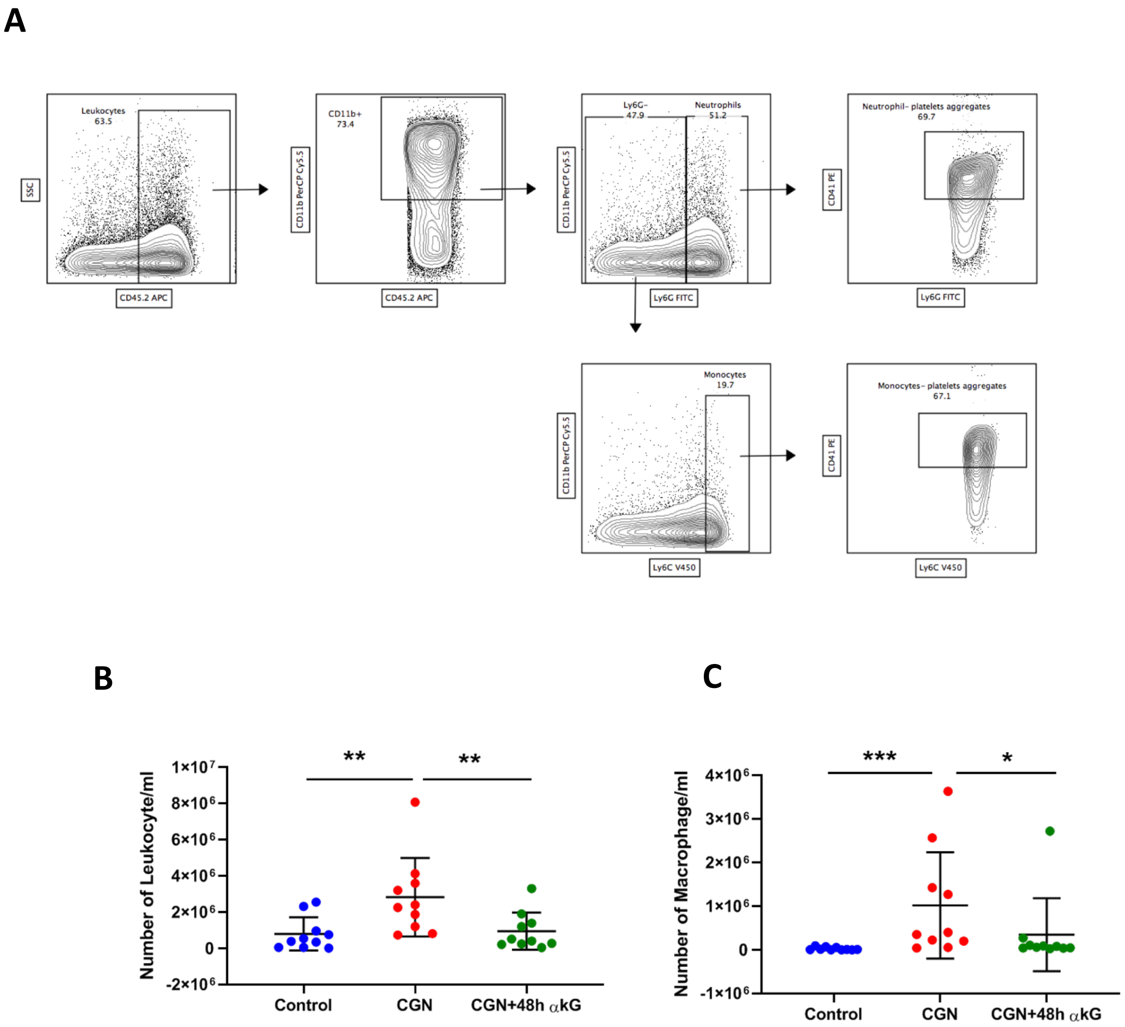


**Supplementary Figure S12. Gating strategy for immune cells from mice peritoneal lavage fluid.** Peritoneal lavage fluid collected from carrageenan treated mice was used to determine number of cells using flow cytometry. **(A)** Represents gating strategy used for determining different cell types. **(B)** Number of leukocytes and **(C)** macrophages in Peritoneal lavage fluid. Data are mean ± SEM. Unpaired t-test was used to compare between the groups, **P*<0.05, ***P*<0.01, ****P*<0.001. Additional data for Figure 5N-T.

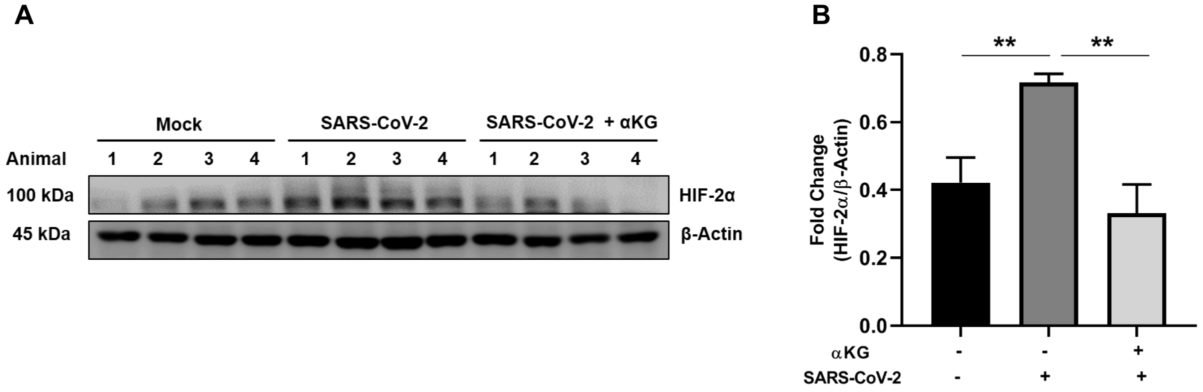


**Supplementary Figure S13. HIF-2α expression in lung tissue of hamsters infected with SARS-CoV-2. (A)** Immunoblots showing reduced expression of HIF2α in lung tissue lysates of hamsters treated with SARS-CoV-2 and supplemented with αKG compared to SARS-CoV-2 treated hamsters. **(B)** Densitometry data of immunoblots from above treatments (n=9 in each group). Data are mean ± SEM. Unpaired t-test was used to compare between the groups, ***P*<0.01.

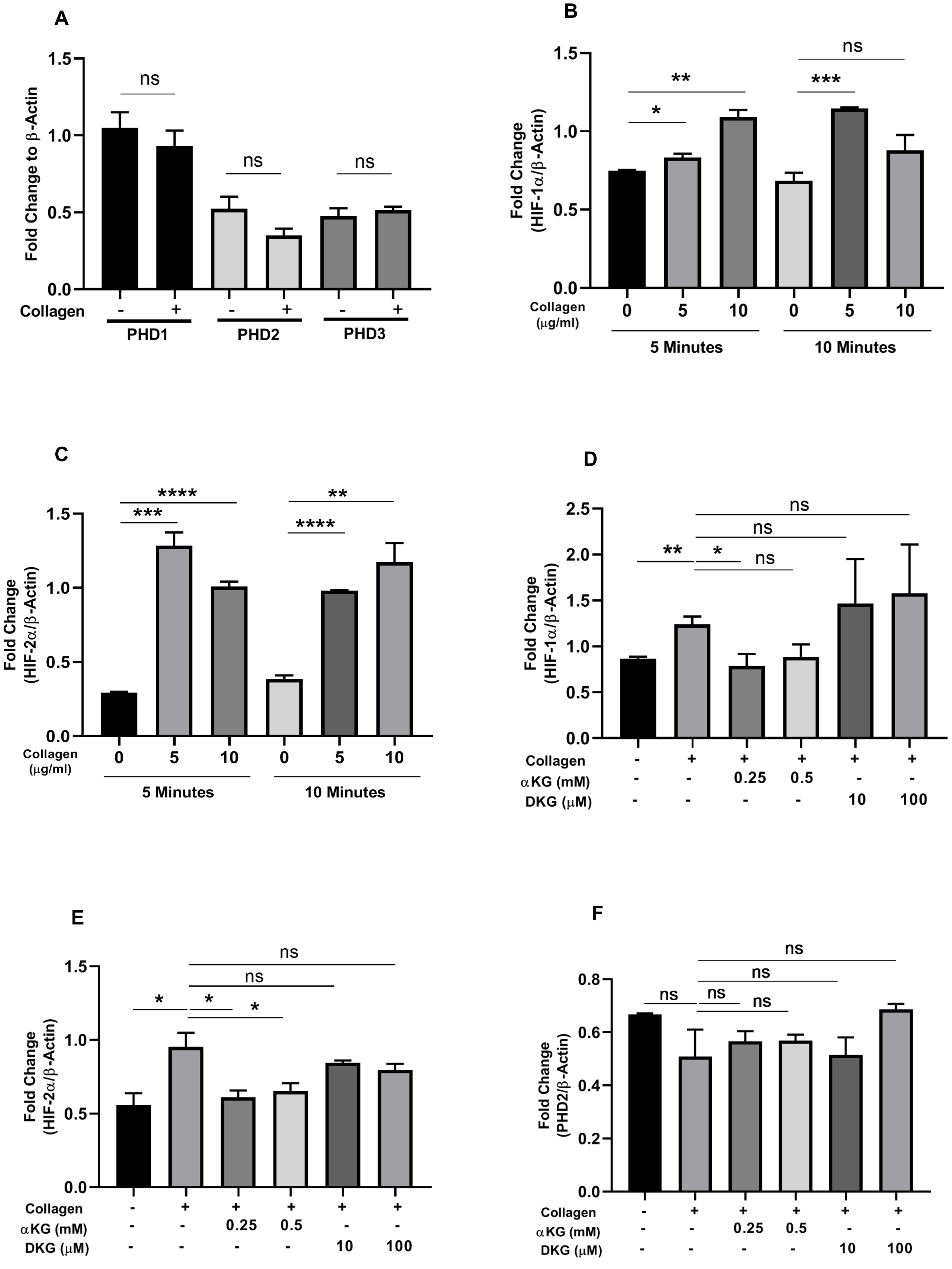
A
Fold Change to β-Actin
1.5
1.0
0.5
0.0
ns
ns
ns
Collagen
- + - + - +
PHD1
PHD2
PHD3
B
Fold Change (HIF-1α/β-Actin)
1.5
1.0
0.5
0.0
*
**
***
ns
Collagen (μg/ml)
0 5 10 0 5 10
5 Minutes
10 Minutes
C
Fold Change (HIF-2α/β-Actin)
1.5
1.0
0.5
0.0
***
****
****
**
Collagen (μg/ml)
0 5 10 0 5 10
5 Minutes
10 Minutes
D
Fold Change (HIF-1α/β-Actin)
2.5
2.0
1.5
1.0
0.5
0.0
**
*
ns
ns
ns
Collagen - + + + + +
αKG (mM) - - 0.25 0.5 - -
DKG (μM) - - - - 10 100
E
Fold Change (HIF-2α/β-Actin)
1.5
1.0
0.5
0.0
*
*
*
ns
ns
Collagen - + + + + +
αKG (mM) - - 0.25 0.5 - -
DKG (μM) - - - - 10 100
F
Fold Change (PHD2/β-Actin)
0.8
0.6
0.4
0.2
0.0
ns
ns
ns
ns
ns
Collagen - + + + + +
αKG (mM) - - 0.25 0.5 - -
DKG (μM) - - - - 10 100

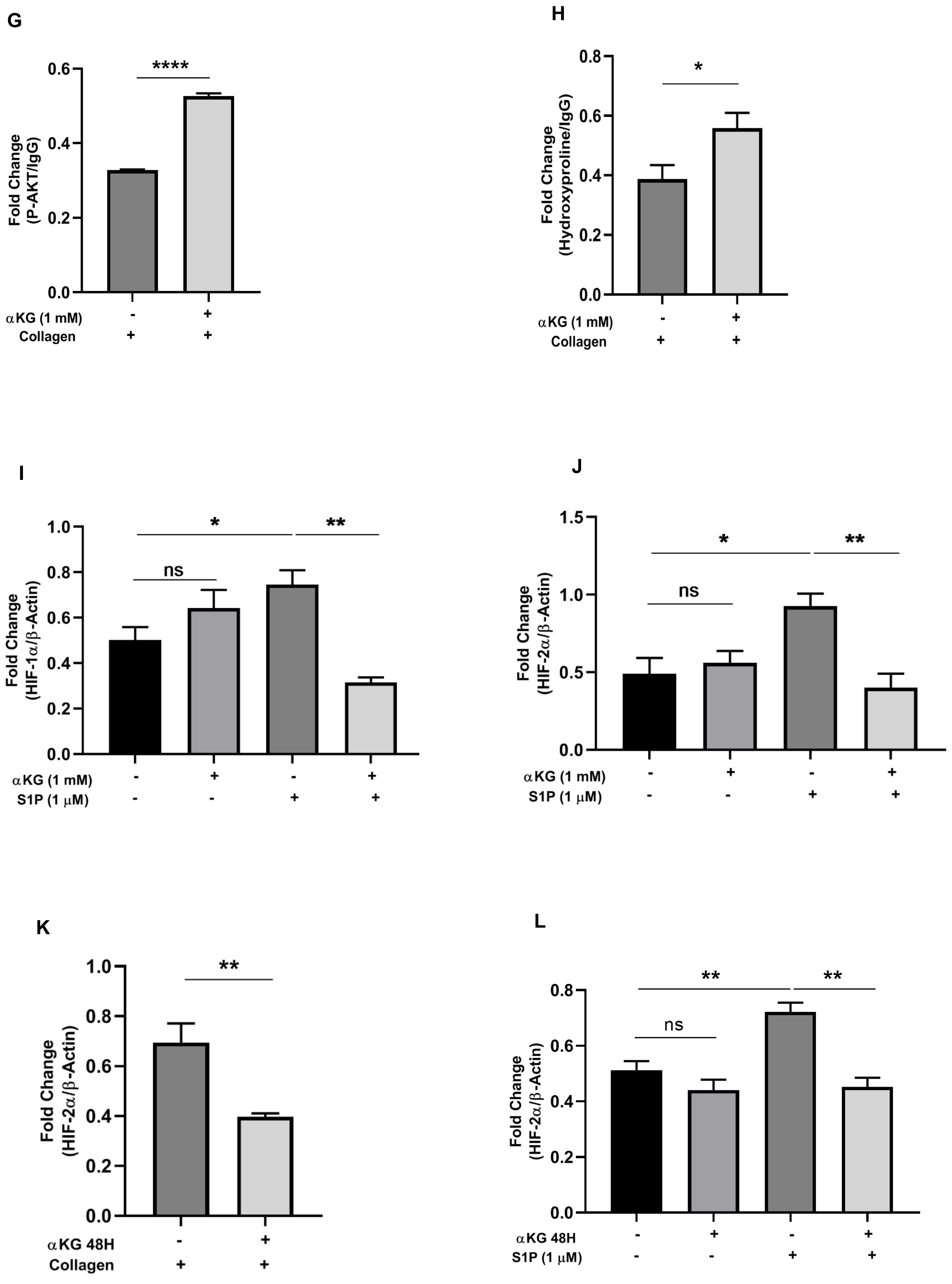
G
****
Fold Change (P-AKT/IgG)
0.0
0.2
0.4
0.6
αKG (1 mM) - +
Collagen + +
H
*
Fold Change (Hydroxyproline/IgG)
0.0
0.2
0.4
0.6
0.8
αKG (1 mM) - +
Collagen + +
I
*
**
ns
Fold Change (HIF-1α/β-Actin)
0.0
0.2
0.4
0.6
0.8
1.0
αKG (1 mM) - + - +
S1P (1 μM) - - + +
J
*
**
ns
Fold Change (HIF-2α/β-Actin)
0.0
0.5
1.0
1.5
αKG (1 mM) - + - +
S1P (1 μM) - - + +
K
**
Fold Change (HIF-2α/β-Actin)
0.0
0.2
0.4
0.6
0.8
1.0
αKG 48H - +
Collagen + +
L
**
**
ns
Fold Change (HIF-2α/β-Actin)
0.0
0.2
0.4
0.6
0.8
αKG 48H - + - +
S1P (1 μM) - - + +

M

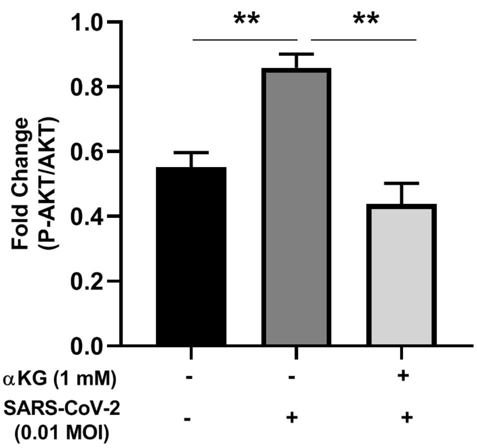

Fold Change (P-AKT/AKT)
1.0
0.8
0.6
0.4
0.2
0.0
**
**
αKG (1 mM) - - +
SARS-CoV-2 (0.01 MOI) - + +


N

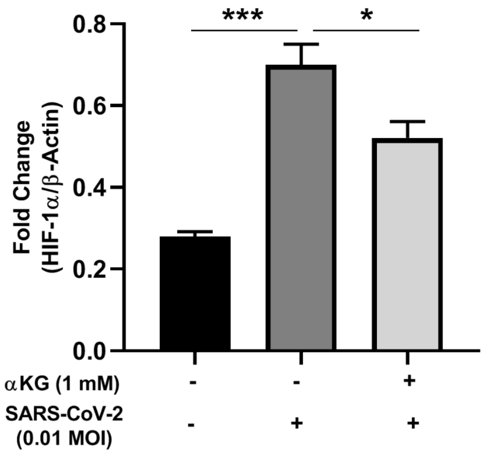

Fold Change (HIF-1α/β-Actin)
0.8
0.6
0.4
0.2
0.0
***
*
αKG (1 mM) - - +
SARS-CoV-2 (0.01 MOI) - + +


O

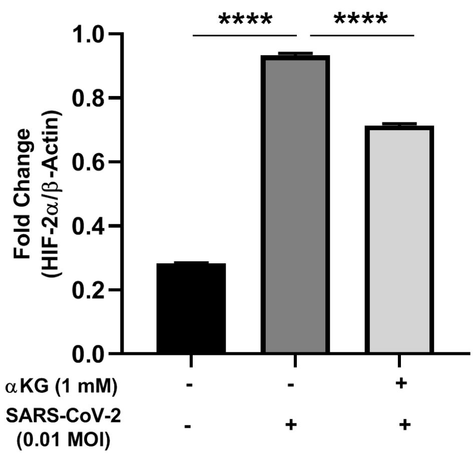

Fold Change (HIF-2α/β-Actin)
1.0
0.8
0.6
0.4
0.2
0.0
****
****
αKG (1 mM) - - +
SARS-CoV-2 (0.01 MOI) - + +


P

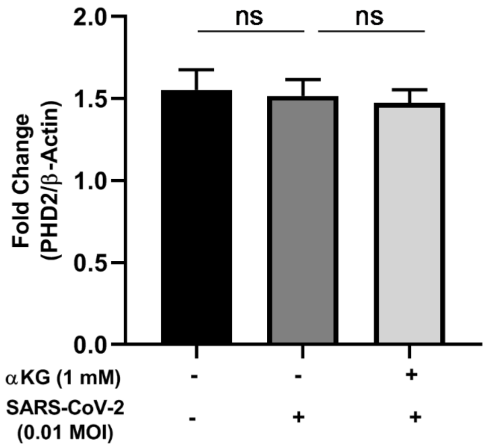

Fold Change (PHD2/β-Actin)
2.0
1.5
1.0
0.5
0.0
ns
ns
αKG (1 mM) - - +
SARS-CoV-2 (0.01 MOI) - + +


Q

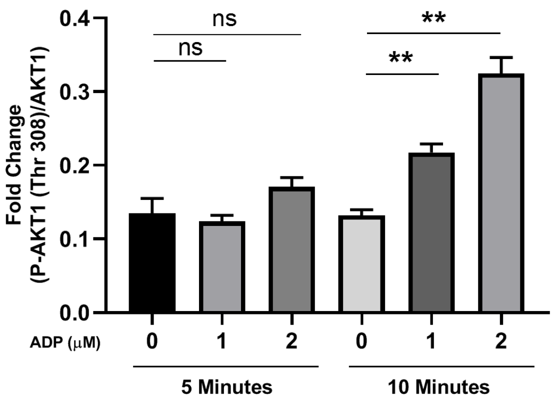

Fold Change (P-AKT1 (Thr 308)/AKT1)
0.4
0.3
0.2
0.1
0.0
ns
ns
**
**
ADP (μM) 0 1 2 0 1 2
5 Minutes
10 Minutes


R

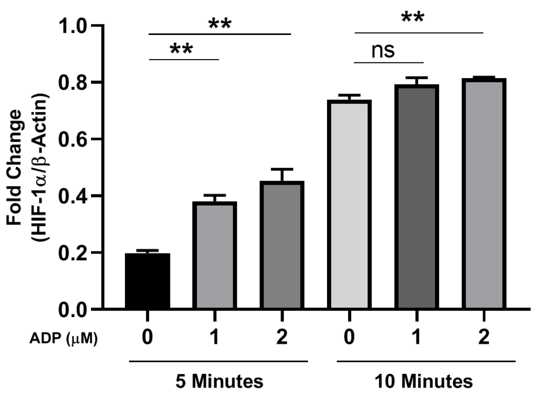

Fold Change (HIF-1α/β-Actin)
1.0
0.8
0.6
0.4
0.2
0.0
**
**
ns
**
ADP (μM) 0 1 2 0 1 2
5 Minutes
10 Minutes

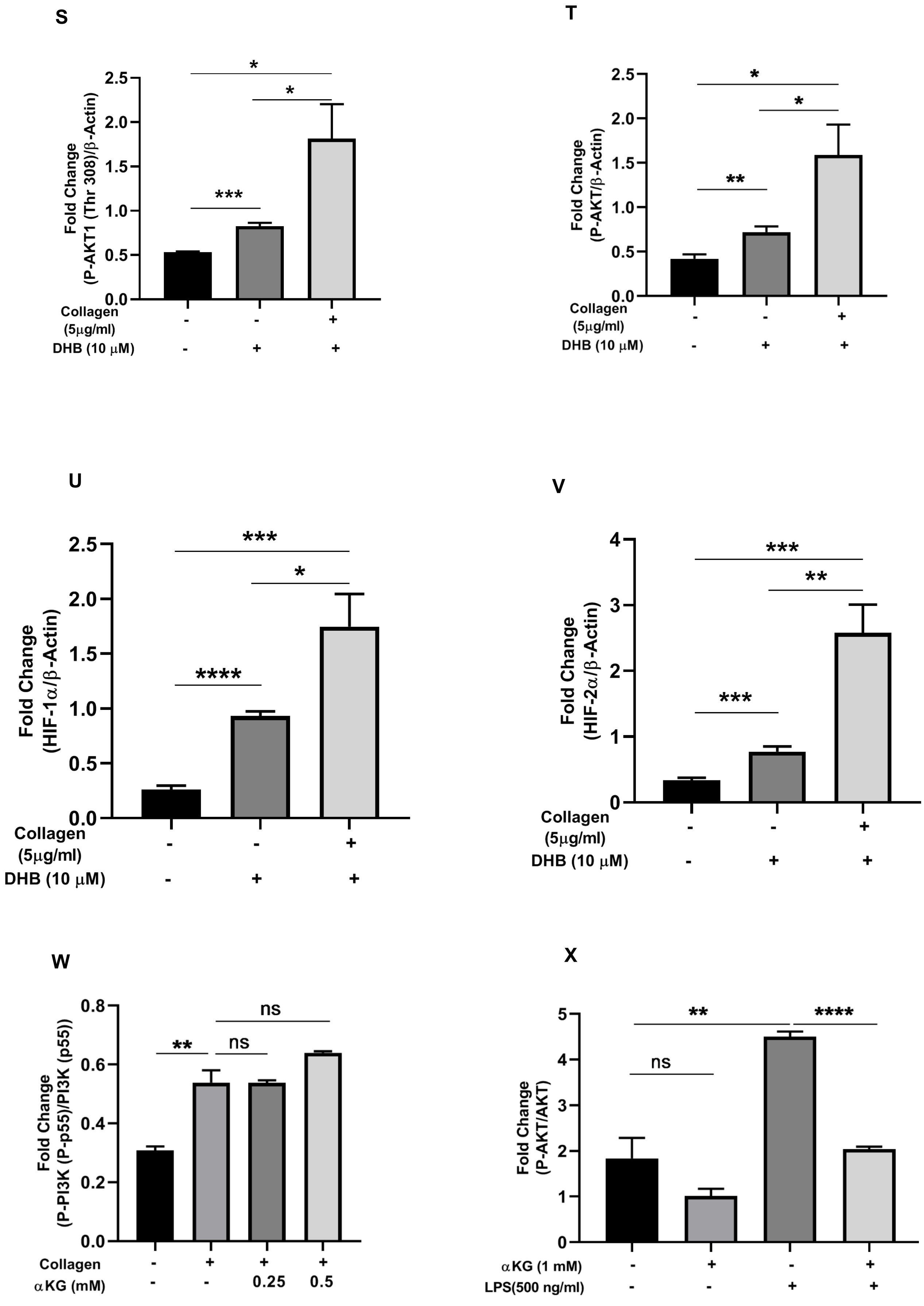
S
Fold Change
(P-AKT1 (Thr 308)/β-Actin)
0.0
0.5
1.0
1.5
2.0
2.5
*
*
***
Collagen
(5μg/ml)
DHB (10 μM)
- - +
- + +
T
Fold Change
(P-AKT/β-Actin)
0.0
0.5
1.0
1.5
2.0
2.5
*
*
**
Collagen
(5μg/ml)
DHB (10 μM)
- - +
- + +
U
Fold Change
(HIF-1α/β-Actin)
0.0
0.5
1.0
1.5
2.0
2.5
***
*
****
Collagen
(5μg/ml)
DHB (10 μM)
- - +
- + +
V
Fold Change
(HIF-2α/β-Actin)
0
1
2
3
4
***
**
***
Collagen
(5μg/ml)
DHB (10 μM)
- - +
- + +
W
Fold Change
(P-PI3K (P-p55)/PI3K (p55))
0.0
0.2
0.4
0.6
0.8
ns
**
ns
Collagen
αKG (mM)
- + + +
- - 0.25 0.5
X
Fold Change
(P-AKT/AKT)
0
1
2
3
4
5
**
****
ns
αKG (1 mM)
LPS(500 ng/ml)
- + - +
- - + +

Y

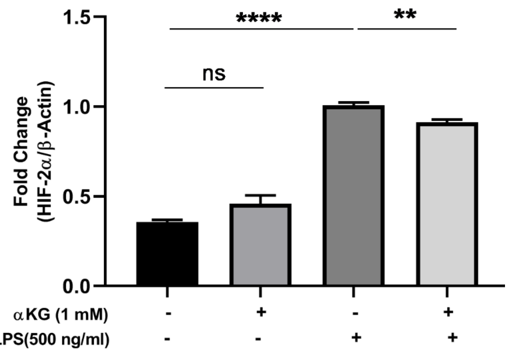


**Supplementary Figure S14. Densitometry of western blots.** Densitometric evaluation of the protein bands was performed using the Alpha imager software for western blots. Densitometry related to western depicted in main Fig. 1A-H (suppl. Fig. 14A-H), Fig. 2A (suppl. Fig. 14I-J), Fig. 4H (suppl. Fig. 14K), Fig. 4N (suppl. Fig. 14L), Fig. 6A (suppl. Fig. 14M-P). Densitometry related to western depicted in suppl. Fig. 1A (suppl. Fig. 14Q-R), suppl. Fig. 1B (suppl. Fig. 14S-V), suppl. Fig. 1C (suppl. Fig. 14W), suppl. Fig. 5 (suppl. Fig. 14X-Y). Bar represents mean ± SEM from triplicate blots. Unpaired t-test was used to compare between the groups, *$P$<0.05, **$P$<0.01, ***$P$<0.001, **** $P$<0.0001, ns=non-significant.

**Supplementary Table S1: List of antibodies used in the study**

| Antibody | Manufacturer | Clone No. | Catalog No. |
|---|---|---|---|
| hu PAC-1 FITC | BD Biosciences | PAC-1 | 340507 |
| hu P-Selectin PE Cy5 | BD Biosciences | AK-4 | 551142 |
| hu Annexin V FITC | BD Biosciences | | 556419 |
| hu CD 41a PE | BD Biosciences | HIP8 | 555467 |
| hu PHD1 | Novus | | NB100-310 |
| hu PHD2 | Novus | | NB100-137 |
| hu PHD3 | Novus | | NB100-139 |
| hu PHD2 | Cell Signalling Technology | | 3293S |
| hu PHD2 | Cell Signalling Technology | D31E11 | 4835S |
| hu HIF1α | Cell Signalling Technology | D2U3T | 14179S |
| hu HIF2α | Invitrogen | ep190b | MA1-16519 |
| hu Phospho PI3K | Cell Signalling Technology | | 4228 |
| hu PI3K | Cell Signalling Technology | | 4292 |
| hu Phospho Akt1 (Thr 308) | Invitrogen | B18H12L21 | 701052 |
| hu AKT1 | Invitrogen | 9Q7 | AHO1112 |
| hu Phospho Akt (PAN) | Cell Signalling Technology | | 9271 |
| hu AKT (PAN) | Cell Signalling Technology | | 9272 |
| Anti-Hydroxyproline (PAN) | Abcam | | ab37067 |
| hu α-tubulin | Invitrogen | TU-01 | 13-8000 |
| hu β-Actin | Cell Signalling Technology | 1300000 | 4970 |
| m CD45.2 APC | Biolegend | 104 | 109814 |
| m CD11b Percp 5.5 | Invitrogen | M1/70 | 45-0112-82 |
| m Ly6C V450 | Invitrogen | HK1.4 | 48-5932-82 |
| m Ly6G FITC | Biolegend | 1A8 | 127606 |
| m CD41a PE | Invitrogen | eBioMWReg30 (MWReg30) | 12-0411-82 |
| m CD11c APC Cy7 | Biolegend | N418 | 117324 |
| mF4/80 PE | Invitrogen | BM8 | 12-4801-82 |
| m HIF1α | Cell Signalling Technology | D2U3T | 14179S |
| m HIF2α | Invitrogen | ep190b | MA1-16519 |
| m Phospho Akt1 (Thr 308) | Invitrogen | B18H12L21 | 701052 |
| m AKT1 | Invitrogen | 9Q7 | AHO1112 |
| m Phospho Akt (PAN) | Cell Signalling Technology | | 9271 |
| m AKT (PAN) | Cell Signalling Technology | | 9272 |
| m α-tubulin | Invitrogen | TU-01 | 13-8000 |
| m β-Actin | Cell Signalling Technology | 1300000 | 4970S |
| Secondary Goat Anti-Rabbit | Cell Signalling Technology | | 7074 |
| Secondary Horse Anti-Mouse | Cell Signalling Technology | | 7076 |
